# Modulating activity of the APC/C regulator SISAMBA improves the sugar and antioxidant content of tomato fruits

**DOI:** 10.1101/2024.11.27.625495

**Authors:** Perla Novais de Oliveira, Leonardo Perez de Souza, Pedro Boscariol Ferreira, Jean Phillippe Mauxion, Luis Felipe Correa da Silva, Ana Isabela Chang, Marina de Lyra Soriano Saleme, Sara Selma García, Herman De Beukelaer, Marina C. M. Martins, Lilian Ellen Pino, Norbert Bollier, Lázaro Eustáquio Pereira Peres, Klaas Vandepoele, Alain Goossens, Nathalie Gonzalez, Alisdair R. Fernie, Nubia Barbosa Eloy

## Abstract

The Anaphase-Promoting Complex/Cyclosome (APC/C) is an E3 ubiquitin ligase that plays a crucial role in ubiquitin-dependent proteolysis of key cell cycle regulators, which is completed by 26S proteasome. Previously, SAMBA, a plant-specific regulator of the APC/C, was identified in Arabidopsis as a critical factor controlling organ size through the regulation of cell proliferation. Here, by assessing its role in the crop tomato (*Solanum lycopersicum*), we confirm that SAMBA is a conserved APC/C regulator in plants. Two *slsamba* genome-edited lines were characterized showing delayed growth, reduced plant size and altered fruit morphology, which were linked to changes in cell division and expansion. Notably, untargeted metabolomics revealed altered flavonoid profiles, along with elevated Brix values in the fruits, indicating a sweeter taste. Accordingly, transcriptomics uncovered a change in temporal gene expression gradients during early fruit development, correlating with the alterations in sugar metabolism and revealing changes in cell wall biosynthesis genes. This study provides the first evidence of *SAMBA’s* role in regulating fruit development, metabolic content, and, ultimately, quality. These important findings offer potential applications for improving the nutritional quality and overall performance of tomato.

## INTRODUCTION

Cell cycle progression must be precisely regulated to achieve proper plant growth, organ development, and reproduction. This process requires the coordinated destruction of essential cell cycle regulatory proteins by specific E3 ubiquitin ligases, known as Anaphase-Promoting Complex/Cyclosome (APC/C) and SKP1-Cullin1-F-box (SCF) complex that recognize proteins to be polyubiquitinated and subjected to proteolysis by the 26S proteasome. The SCF complex is crucial at the G1 to S phase, where it targets cell cycle-dependent kinase inhibitors (CKIs), including Substrate/Subunit Inhibitor of Cyclin-dependent protein kinase (SIC1) in yeast (Dirick et al., 1995) and Kip-related proteins (KRPs) in plants (Verkest et al., 2005; Ren et al., 2008). In contrast, the APC/C complex primarily functions at the G2 to M transition and mitotic exit. For example, to transition from G2 to M phase and exit mitosis, mitotic cyclins and Securin must be targeted for degradation by the APC/C in all organisms (Petersen et al., 2000; Harper et al., 2002; Capron et al., 2003; Buschhorn and Peters, 2006). Although the functions of these multi-protein machines have been primarily investigated in *Arabidopsis thaliana*, other plant species remain less explored. Hence, understanding its impact on crops such as tomato can reveal unique regulatory mechanisms and potential applications in agriculture and biotechnology, enhancing the translatability of basic research findings for crop improvement (Inzé and Nelissen, 2022).

The APC/C in plants can comprise up to 14 subunits, such as in Arabidopsis, maize, and sorghum, which are divided into at least three main functional modules: a catalytic/substrate recognition module (APC2, APC11, and APC10), a structural module (APC3, APC6, APC7, and APC8) (D’Andrea and Regan, 2003; Alfieri et al., 2017), and a scaffolding module to which the catalytic and structural components are attached (APC1, APC4, and APC5) (Thornton and Toczyski, 2003; Thornton et al., 2006; Eloy et al., 2015; Alfieri et al., 2017). Moreover, the APC/C activity is regulated by two structurally related co-activator proteins, CELL DIVISION CYCLE 20 (CDC20) and CELL CYCLE SWITCH 52 (CCS52) (Peters, 2002; Baker et al., 2007), and inhibited by ULTRAVIOLET-B-INSENSITIVE4 (UVI4) and its homolog OMISSION OF SECOND DIVISION 1 (OSD1)/GIGAS CELL 1 (GIGAS)/ UVI4-Like (d’Erfurth et al., 2009; Heyman et al., 2011; Iwata et al., 2011).

Functional characterization of APC/C subunits in Arabidopsis has revealed their essential roles in cell differentiation, development of the shoot and root meristems, plant growth, vascular development, hormone regulation, and endoreduplication (Blilou et al. 2002; Saze and Kakutani, 2007; Marrocco et al., 2009; Rojas et al., 2009; Eloy et al., 2011; Eloy et al. 2012; Schwedersky et al., 2021). Moreover, in *Medicago truncatula*, the APC6 regulates the number of lateral roots and nodule formation (Kuppusamy et al., 2009). In *Oryza sativa*, APC6 knockout plants show reduced height and smaller cell size, while the TE gene, a homolog of CCS52A, controls shoot branching and tillering (Kumar et al., 2010; Lin et al., 2012).

Some years ago, we identified a plant-specific APC/C regulator named SAMBA (Eloy et al., 2012). Loss-of-function mutation of *SAMBA* in Arabidopsis led to increased cell proliferation, resulting in larger leaves, roots, and seeds. In maize, Clustered Regularly Interspaced Palindromic Repeats/CRISPR-associated protein 9 (CRISPR/Cas9) *zmsamba* mutants also show an increased rate of cell division. However, these plants displayed reduced organ and tissue growth, resulting in dwarfism as a consequence of decreased cell size (Gong et al., 2021). In addition to these contrasting phenotypes, the *SAMBA* expression pattern also varies between these species during development. In Arabidopsis, *SAMBA* is highly expressed during embryogenesis, with transcripts gradually decreasing when seedlings germinate, being restricted to the hypocotyl at 8 days after stratification (DAS), and exclusively detected in pollen grains at more advanced developmental stages. By contrast, *ZmSAMBA* expression in maize is more constant throughout the entire development (Sekhon et al., 2011).

Here, we identified and investigated the role of *SlSAMBA* in tomato plants, an excellent model for studying fleshy fruit development. Tomato is particularly interesting due to its significant economic value, short life cycle, the availability of extensive genomic resources, and the established protocols for genetic manipulation (Zhang et al. 2016). After pollination, tomato fruits undergo a very orchestrated growth journey, progressing through successive and overlapping phases of cell division, expansion, and ripening (Mauxion et al, 2021; Mumtaz et al. 2022). This growth period, during which cell proliferation, cell expansion, and endoreduplication will determine final fruit size, is accompanied by important metabolic changes leading to the formation of mature tomato fruits rich in structurally diverse metabolites. The consumption of tomato has been associated with health benefits including a lower incidence of several chronic diseases and certain types of cancer (Giovannucci, 1999; Willcox et al., 2003), which are often attributed to the high levels of antioxidant secondary compounds, particularly flavonoids, that accumulate in the fruit (Zhang et al., 2015; Martin and Li, 2017).

To gain a deeper understanding of *SlSAMBA’*s functional role, we employed CRISPR/Cas9-based genome editing to generate *SlSAMBA* loss-of-function plants (*slsamba*) in the Micro-Tom tomato. These mutants exhibited delayed growth and altered development demonstrated by reductions in both plant size and fruit dimensions. Alterations in fruit morphology, particularly a shift toward elongated shapes and impaired seed set, suggest that SlSAMBA plays a critical role in regulating pistil/ovary development, as well as reproductive success in tomato. Moreover, metabolomic profiling identified higher contents of soluble sugars and compounds related to flavonoids in the *slsamba* fruits. These combined results show that SISAMBA plays a key role in regulating various aspects of tomato fruit development, with potential implications for improving crop quality.

## RESULTS

### The tomato genome contains the SAMBA gene highly expressed in the early stages of flower and fruit development

The Arabidopsis SAMBA (AT1G32310) protein contains three putative motifs (SHR1, LCR, and SHR2) previously identified by Eloy et al. (2012), which appear to be conserved across most plant species (Figure 1A). Our search for a SAMBA homolog in tomato identified a single gene, *Solyc08g076580.2* (hereafter referred to as *SISAMBA*). The open reading frame of *SISAMBA* has 342 base pairs (bp) and encodes a 113-amino acid protein. Our phylogenetic analysis shows that SlSAMBA protein is closely related to its counterpart in tobacco (Figure 1A) and multiple sequence alignment reveals high similarity to other homologs (Figure 1B). Data from the general feature format (GFF) file provided with the genome sequence revealed that the *SlSAMBA* gene is located at the end of the long arm of chromosome 8.

**Figure 1.**
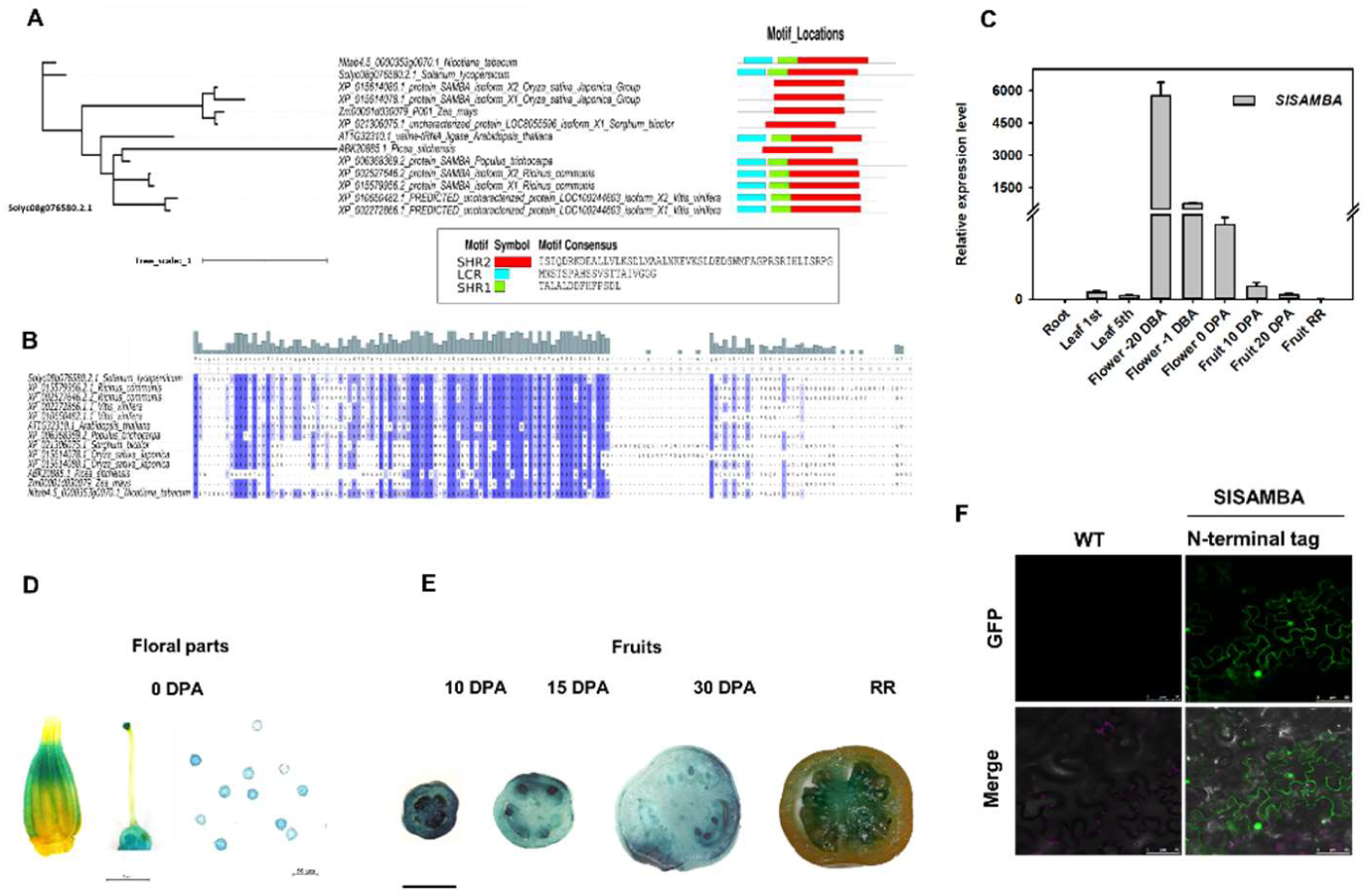
Phylogenetic relationships and architecture of conserved motifs in the SISAMBA protein and spatio-temporal expression of *SAMBA* in tomato cv. Micro-Tom. (A) Phylogenetic tree of SAMBA proteins from *Solanum lycopersicum*, *Ricinus communis*, *Vitis vinifera*, *Arabidopsis thaliana*, *Populus trichocarpa*, *Sorghum bicolor*, *Oryza sativa*, *Picea sitchensis*, *Zea mays*, and *Nicotiana tabacum* constructed using IQ-Tree v.1.6.9. Three putative conserved motifs (SHR1, LCR, and SHR2) described by Eloy et al. (2012) were identified in most SAMBA homologs. Their location and motif consensus sequences are shown. (B) Multiple sequence alignment of the *S. lycopersicum Solyc08g076580.2.1* gene (SAMBA-homolog) was performed using Clustal X version 2.1 with default parameters (Thompson et al., 1997). (C) Tissue-specific expression profile of *SlSAMBA*. Data was generated by qRT-PCR, using *β-ACTIN* as an internal control. Root (30-day-old plants), 1^st^ leaf, and 5^th^ leaf in logarithmic scale, with 26 to 500 on the x-axis omitted; 20 days before anthesis (−20 DBA), −1 DBA, flowers at the pre-anthesis stage, 1 day before anthesis; and 0 DPA, the day of anthesis; 10 DPA, 10 days post anthesis, 20 DPA, 20 days post anthesis; and RR, red ripe fruit. (D) *SISAMBA*-promoter-directed expression of GUS in tomato transformed plants (T3). *SISAMBA* expression was high in anther, pistil, and pollen grains (Scale bar = anther and pistil (1mm), pollen grains (50 µm), and fruits (2 mm). (E) *SISAMBA* expression decreased gradually in later stages of fruit development. (F) SISAMBA:EGFP (N-terminal) tagged protein is localized in the nucleus and spread throughout the cytoplasm. The upper panel shows the GFP signal field, and the lower panel shows the merged fields. Scale bar = 50 µm.

We assessed the mRNA levels of *SlSAMBA* in different organs and developmental stages by quantitative Reverse Transcriptase PCR (qRT-PCR) (Figure 1C). *SlSAMBA* showed the highest expression in floral buds (20 days before anthesis (−20 DBA) and −1 DBA, which decreased in flowers at anthesis (0 days post anthesis - DPA). In the fruit, *SlSAMBA* was more expressed at 10 DPA compared to 20 DPA and the red ripe stage (52 DPA). In 30-day-old plants, *SISAMBA* transcripts were undetectable in the root, but were present in the leaves, with higher levels in the early (first leaf) than in the later (fifth leaf) stages of development. Additionally, in silico analysis of public transcriptome data from Plant eFP (Waese et al., 2017) revealed that *SlSAMBA* is widely expressed in all organs (Figure S1), with higher expression levels during early developmental stages. This finding supports our qRT-PCR data and is consistent with the essential role of SISAMBA in cell cycle regulation.

To further analyze the expression pattern of *SlSAMBA*, a 1.6-kb fragment upstream of its ATG start codon was cloned into the pKGWFS7 vector to drive the expression of β-glucuronidase/Green Fluorescent Protein (GUS-GFP) reporters, and the construct was introduced into Micro-Tom tomato plants (Figure S2A). *SlSAMBA* expression was particularly high during embryogenesis (Figure S2B) and was observed in different parts of the pistil, including the ovary and stigma, but not in the style. In the androecium, *SlSAMBA* was expressed in both the anther and pollen grains, as shown in Figure 1D. In later stages of fruit development, the highest *SlSAMBA* expression occurred at 10 DPA, a phase known for intense cell division, and decreased afterward until the ripening stage, when the expression was restricted to the septum and columella (Figure 1E). To determine the subcellular localization of SlSAMBA, we fused its N-terminal region with eGFP (Figure 1F; Figure S2C). The eGFP-SISAMBA fusion was transiently expressed in epidermal cells of 30-day-old tobacco leaves through Agrobacterium infiltration. The green fluorescence emission was detected in the cytoplasm and nuclei, confirming the *in silico* analyses (Table S1 and S2).

### SlSAMBA interacts with APC6 and APC3b, two members of the APC/C complex

To identify protein interactors of SlSAMBA, we employed TurboID-Mediated Proximity Labeling in tomato hairy roots (Arora et al. 2020). This approach is based on the *in planta* expression of the promiscuous biotin ligase TurboID fused to a target protein of interest, after which streptavidin-based affinity purification is used to capture the biotinylated proteins that (in)directly interact with or sit in proximity to the target protein. An advantage of the TurboID system is its ability to detect indirect interactions, which is particularly useful for studying large protein complexes (Zhang et al., 2022). The SlSAMBA-TurboID fusion construct was stably expressed in tomato hairy roots and compared to controls to verify intact protein expression and (auto-) biotinylation activity of the TurboID-fusion proteins.

A total of 34 (at a permissive false discovery rate (FDR) of 0.01) or 16 (at a stringent FDR of 0.001) proteins were significantly enriched in the SlSAMBA-TurboID samples compared to the eGFP-TurboID controls (Dataset S1). Seven out of the 16 exhibited over four-fold enrichment (log_2_) among these interacting or proximal proteins. Notably, two of the interactors were identified as tomato APC/C subunits APC6 and APC3b (Dataset S1, Figure S3A-B), and another two as homologs of Arabidopsis NAP1-RELATED PROTEIN 2 (NRP2) involved in the cell cycle. This demonstrates that, as in other plant species, SISAMBA is a member of APC/C complex in tomato (Eloy et al., 2012).

To confirm the Turbo ID results, which showed that SISAMBA interacts with two core APC/C subunits in tomato, we performed a yeast two-hybrid (Y2H) assay. We tested the interaction of SISAMBA with SIAPC3b and SIAPC10 subunits. Our results showed a direct interaction between SISAMBA and SIAPC3b, but no interaction with SIAPC10 (Figure S3C), which was consistent with our previous findings in Arabidopsis (Eloy et al, 2012).

### SISAMBA frameshift mutants exhibit a dwarf phenotype with reduced organ size

We next used CRISPR/Cas9-based gene editing to induce mutations in *SlSAMBA* and characterize the phenotype of the edited plants. To generate CRISPR-Cas9 *slsamba* mutants, a dual gRNA approach was used, employing two distinct gRNA combinations to transform tomato (Figure S4A). After sequencing 65 selected transformants, we identified two plants showing different mutations. The *slsamba*-27 mutant (construct 1) features a single nucleotide deletion at the first target site (gRNA3), resulting in a stop codon at position 178-180 and potentially an in-frame deletion of 53 amino acids (Figure S4B, C). The *slsamba*-3 mutant (construct 2) carries a 73-nucleotide deletion between the two gRNAs (Figure S4B), resulting in a premature stop codon and potentially a shorter protein of 61 amino acids (Figure S4C). Alternatively, if the aberrant mRNA is degraded, no protein will be produced.

The two T0 *slsamba* mutants were backcrossed with wild-type (WT) and, later, T1 heterozygous plants without the Cas9 sequence were selected and self-pollinated. In the fourth generation (T4), we obtained homozygous mutants and qRT-PCR analysis of whole plants confirmed that both *slsamba* lines have very low relative expression levels of *SlSAMBA* compared to the WT (Figure S4D).

To assess the impact of *SlSAMBA* mutation on tomato development, we measured several growth-related parameters at both the vegetative and reproductive stages. The edited plants were dwarf (Figure 2A), with the height of the first inflorescence significantly shorter by 56.2 % (line 3) and 46.9% (line 27) compared to the WT (Figure 2B). Additionally, they showed reduced stem diameter (Figure 2C) and smaller leaf area (Figure 2D-E).

**Figure 2.**
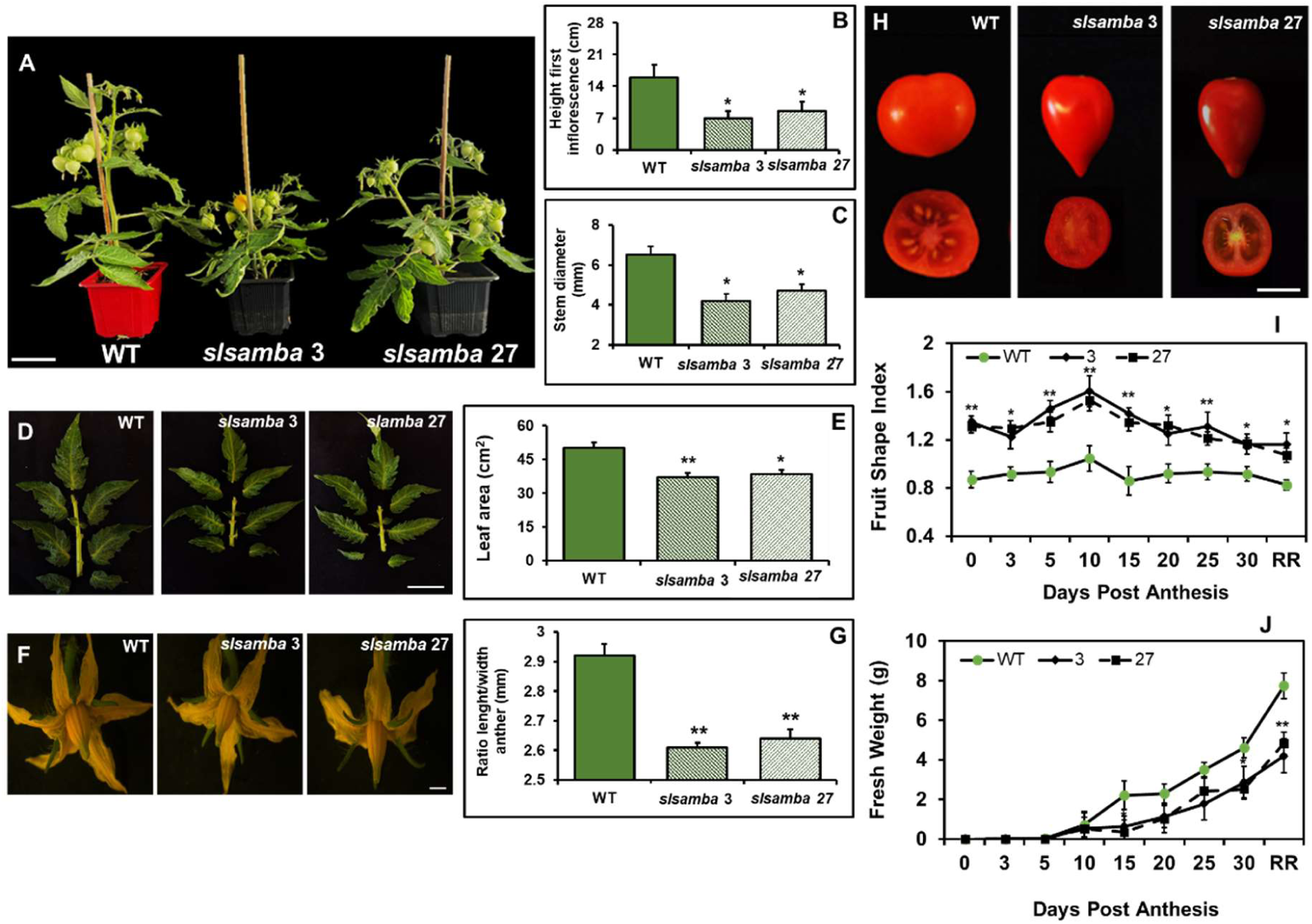
Phenotypic effects of *samba* edition on tomato plant development. (A) Representative 40-day-old *slsamba* (#3 and #27) and WT plants grown in soil. The edited lines are smaller compared to the WT. (B-C) Height of the first inflorescence and stem diameter of WT and *slsamba* mutants. Data are means ± SEM (n = 12). (D-E) Morphology of 5^th^ leaf in WT and *slsamba* and total leaflet area of one leaf measured 1 month after sowing. Data are means ± SEM (n = 21). (F-G) Flowers at anthesis in WT and *slsamba* were observed by stereo microscopy (SMZ 1500 increased 7.5×). *slsamba* mutants have a reduced average length/width ratio anther at anthesis. Data are means ± SEM (*n* = 21). (H) Mature fruits of WT and *slsamba* plants. (I-K) Fruit shape index and weight of the third fruit per inflorescence for WT and *slsamba* at different stages. The disruption of *SISAMBA* function altered fruit weight and shape and reduced their average length/width ratio. Data are means ± SEM (*n* = 24). Significant differences (ANOVA followed by Dunnett’s test) are indicated by asterisks (*P < 0.05 and **P < 0.01).

Since Arabidopsis *SAMBA* mutants exhibit defects in male gametophyte development (Eloy et al., 2012), we investigated whether similar phenotypic alterations were present in our tomato-edited lines. The *SISAMBA* mutants produced smaller flowers with thinner anthers at anthesis (Figure 2F-G). Furthermore, their mature fruits were more elongated and thinner compared to WT (Figure 2H-I), with reduced weight throughout development (Figure 2I-J). By measuring the fruit shape index, we found that the elongation phenotype was evident as early as anthesis, and it was linked to a decrease in both diameter and weight.

### Effect of SlSAMBA mutation on female and male gametophytes

We observed that fruits from homozygous *slsamba* plants contained notably fewer viable seeds, with an average of 2.1 seeds per fruit (∼8% of the average 26.1 from the WT) (Figure 3A). The reduced seed viability suggests potential issues in gametogenesis. To explore this further, we analyzed the phenotypes of the flowers in detail, with a particular focus on the ovary and pollen.

**Figure 3.**
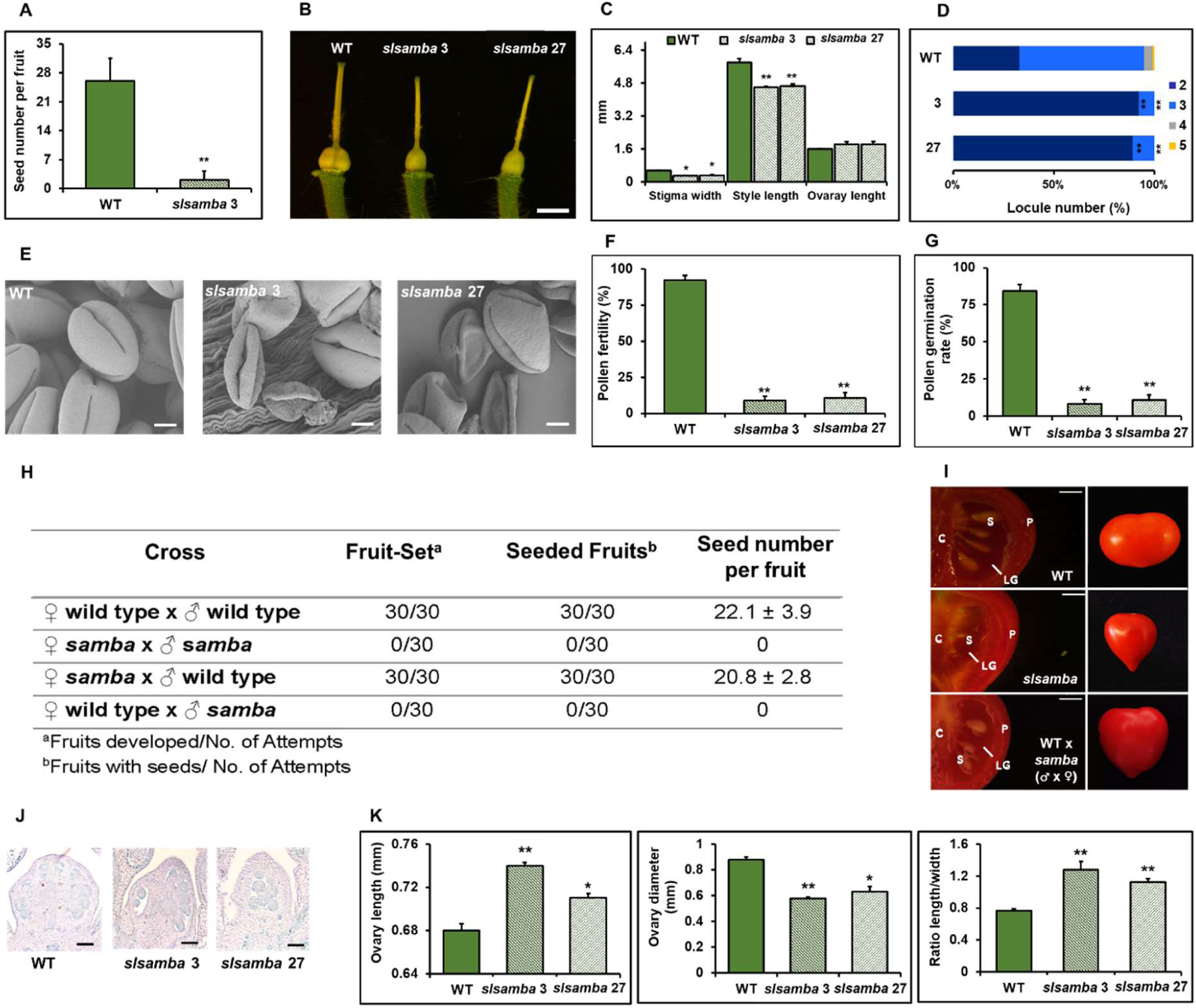
Phenotypic effects of *slsamba* mutation on female and male gametophytes. (A) Seed number per fruit from spontaneous self-pollinations (♀WT × ♂WT/ ♀*slsamba-3* × ♂*slsamba-3*). (B-C) Pistil phenotypes of WT and *slsamba* plants; Scale bar = 2 mm. The edited lines have shorter stigma width and style length compared to the WT, but the ovary length is not statistically different. Data are means ± SEM. (*n* = 21). (D) Stacked bar plots showing the percentages of fruits with different numbers of locules in WT and *slsamba* mutants (n = 8 plants). (E) Electron micrographs of WT and *slsamba* pollen grains (Scale bar = 5 µm). (F-G) Pollen fertility [(number of viable pollen grains)/(number of total pollen grains counted × 100)] and pollen germination [(number of germinated pollen grains)/(number of total pollen grains counted × 100)]. Data are means ± SE considering 10 optical fields randomly selected. (H) Total fruit set and seeded fruits from WT or *slsamba* emasculated flowers crosses (♀WT × ♂WT/♀*slsamba* × ♂*slsamba*) and back-crosses (♀*slsamba* × ♂WT/ ♀WT × ♂*slsamba*). (I) Representative longitudinal and whole sections of WT, *slsamba*, and ♂WT x ♀*slsamba* ripe fruits, where seeds can be observed more frequently in WT and ♂WT x ♀*slsamba* but not in the mutant. C, columella; S, seed; P, pericarp; LG, locular gel; Scale bar = 2 mm. (J) Histological analysis of longitudinal sections of pistils at −6 days before anthesis (DBA) of WT and *slsamba* mutants; Bars = 100 µM. (K). The average ovary diameter, ovary length, and ovary shape index were calculated using ImageJ. Data are means ± SEM (n = 9). Significant differences (ANOVA followed by Dunnett’s t test) are shown by asterisks (*P < 0.05 and **P < 0.01).

The ovary morphology of *slsamba* mutants (lines 3 and 27) exhibited shorter stigma width and style length compared to WT plants at anthesis; however, ovary length did not show a significant difference in this stage (Figures 3B-C). Additionally, locule number quantification revealed that over 90% of the fruits from both edited lines contained two locules, whereas the WT predominantly had three locules (Figure 3D). Pollen analysis showed low fertility and germination rates in *slsamba* mutants, with only 9.5% (line 3) and 11.3% (line 27) of anthers being fertile, and 9.7% (line 3) and 12.8% (line 27) of pollen grains germinating. These results were partially explained by the aberrant morphology of pollen grains from the edited lines compared to WT pollen, as illustrated in the electron microscopy images shown in Figure 3E–G.

To pinpoint the effect of *SISAMBA* knockout on fruit phenotype, specifically fruit shape and seed formation, we conducted reciprocal crosses between *slsamba-3* x WT (♀*slsamba* ×♂WT/ ♀WT × ♂*slsamba*) (Figure 3H-I). When heterozygous *slsamba* mutants were used as pollen recipients, the altered fruit shape was maintained, suggesting a persistent effect on fruit morphology, likely resulting from defective male gametogenesis. This is evident in fruits from back-crosses of emasculated *slsamba* flowers (♀*slsamba-3* × ♂WT), wich produced a substantial number of seeds, while no seeds were formed when WT plants were used as recipients (♀WT × ♂*slsamba-3*).

Since fruits are derived from ovarian tissues after pollination, we investigated whether the elongated shape in *slsamba* mutants is determined at pre-anthesis. The comparison between pistils from *slsamba* and WT at 6 DBA shows that mutants have longer ovaries with reduced diameter, resulting in an increased shape index (Figure 3J-K). These results indicate that the *SISAMBA* mutation alters ovary morphology before anthesis, likely triggering changes in fruit shape.

The tomato fruit comprises distinct complex tissues, including pericarp (divided into exocarp, mesocarp, and endocarp), placenta, and septum (Mauxion et al., 2021). The pericarp represents about two-thirds of the total fruit weight and plays an important role in determining the fruit’s quality. The development of the tomato pericarp includes an intense phase of cell division closely linked to changes in the cell cycle, followed by cell expansion (Inzé and Veylder, 2006). To determine whether the observed morphological changes in the ovary and fruit shape were associated with alterations in ploidy levels, we performed flow cytometry analysis on nuclei isolated from the pericarp at 10 to 30 DPA. The analysis revealed that ploidy profiles in the *slsamba* mutants were not altered (Figure S5). We investigated the cellular effect of the *SISAMBA* mutation during pericarp development by performing a time course analysis of pericarp growth from anthesis to 30 DPA (pericarp fixed in FAA at 0, 10, 15, and 30 DPA) (Figure 4A). We observed that the fruits of *slsamba* mutants displayed an increased pericarp thickness compared to the WT (Figure 4B). Additionally, by counting the number of cell layers from the epidermis to the endodermis, we observed an increase in cell layers in *slsamba* fruits, suggesting higher cell division activity (Figure 4C). Furthermore, *slsamba* mutants exhibited smaller cell sizes, although this difference diminished as fruit development progressed (Figure 4D). Notably, alterations in the mesocarp of WT and *slsamba* fruits were more evident at 0 DPA and 10 DPA. We also observed that mesocarp cells in *slsamba* mutant fruits were more irregular in shape than those in WT fruits, with a more elongated appearance. Additionally, mutant fruits showed a 70% increase in the number of cells per area along the mediolateral axis compared to WT fruits, indicating a higher cell density within the same area (Figure 4E).

**Figure 4.**
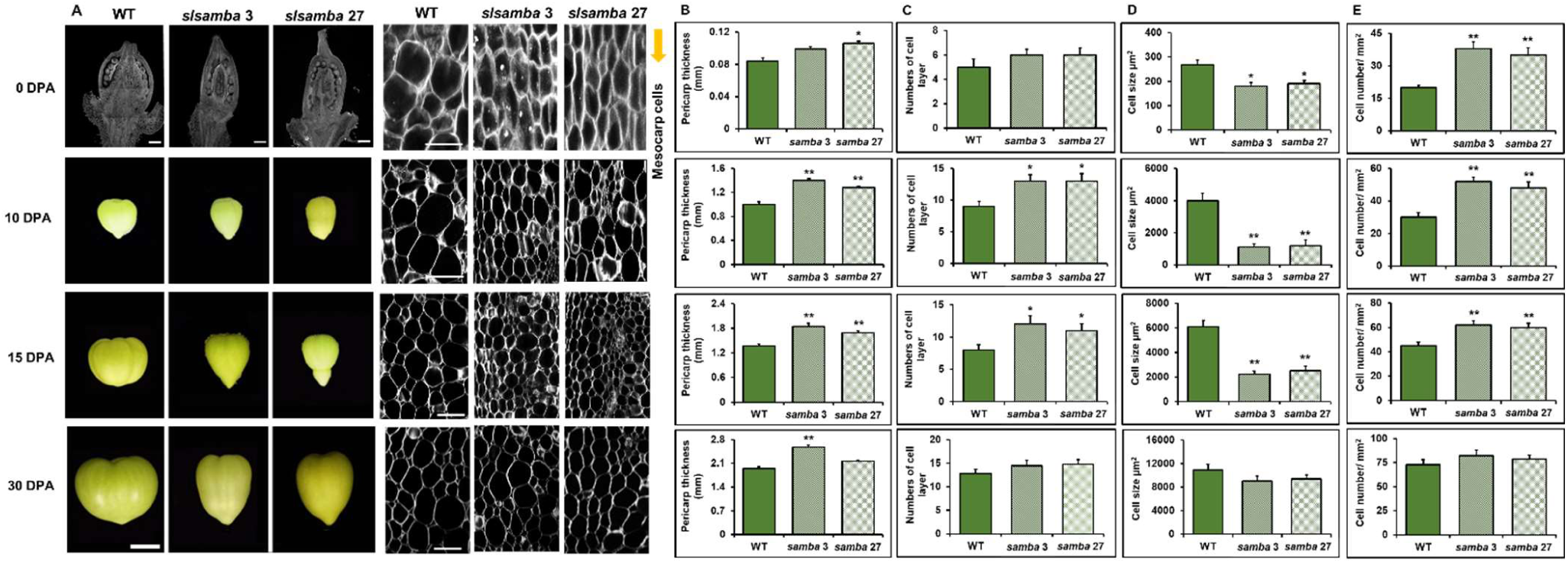
Microscopic observations of WT and *slsamba* fruits. (A) Morphology of fruits and microscopic observations of wild-type Micro-Tom (WT) and *slsamba* plants (#3 and #27) at 0 (Bar = 300 µM), 10 (Bar = 20 µM), 15 (Bar = 20 µM), and 30 days after anthesis (DAP) (Bar = 200 µM). (B) Comparison of pericarp thickness, (C) number of cell layers of the pericarp tissues, (D) average cell size, and (E) cell numbers per area in the mesocarp regions in the WT and *slsamba* mutants. Data are means ± SEM (n = 9). Significant differences (ANOVA followed by Dunnett’s t test) are shown by asterisks (*P < 0.05 and **P < 0.01).

### SlSAMBA mutation broadly alters primary and secondary metabolism

We further explored whether the altered fruit phenotype in the *slsamba*-edited plants is associated with or accompanied by changes in their metabolic profiles. Since *SISAMBA* expression is highest during the early stages of fruit development (Figure 1C), fruit samples were collected at 3-, 5- and 8 DPA, and their primary and specialized metabolite contents were analyzed by gas chromatography coupled with mass spectrometry (GC-MS) and liquid chromatography coupled with mass spectrometry (LC-MS), respectively. We identified 41 primary metabolites including 17 amino acids, 13 organic acids, three sugars, two fatty acids, and two sugar alcohols (Table S3). The heat map representing metabolite accumulation in the different genotypes revealed several differences between *slsamba* and WT that collectively contributed to clustering samples in partial least square-discriminant analysis (PLS-DA) (Figure S6). As expected, samples from both *slsamba* lines were overlapped and separated from the WT on PLS-DA components 1 and 2, which together explained 40.3% of the total variance. The most important metabolites contributing to this separation (variable importance in projection (VIP) > 1.5) were glucose, aspartic acid, sucrose, oxo-glutaric acid, and GABA (Figure S6C).

Given that primary metabolites are major components of fruit quality (Beauvoit et al. 2014), we examined the significant differences in *slsamba* lines compared to WT in more detail (Figure 5). Except for glucose, soluble sugars were usually at higher levels in the fruits of *slsamba* mutants. This result is in agreement with the high sucrose demand during the cell division stage of fruit development (Liu et al., 2007; Biais et al., 2014).

**Figure 5.**
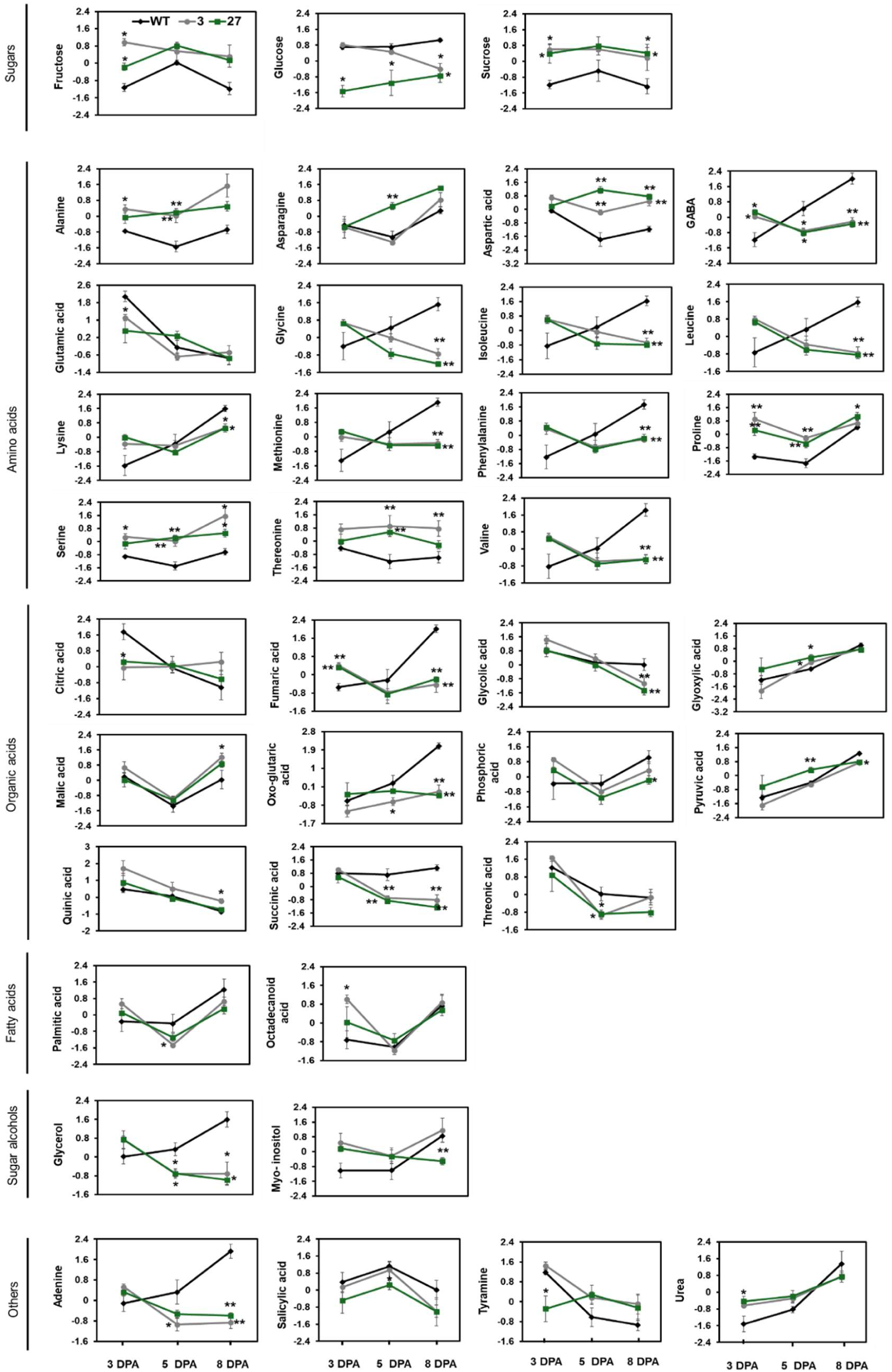
Primary metabolites significantly altered in fruits of *slsamba* mutants compared to the WT. Relative metabolite quantification was performed by gas chromatography coupled with tandem mass spectrometry (GC-MS/MS) using fruits from individual homozygous plants of lines 3 and 27 and WT plants harvested at three developmental stages (3-, 5- and 8-days post anthesis). Metabolites were identified based on receiver operating characteristic (ROC) curves using Log and Auto-scaling normalized data on the MetaboAnalyst platform 6.0. Data are means ± SD (n = 4-6). Gray and green lines indicate lines 3 and 27, respectively. Significant differences (one-way ANOVA, post hoc paired t-test) between the WT and each mutant line are indicated by asterisks (*P < 0.05 and **P < 0.01).

The levels of most amino acids also varied significantly between *slsamba* and WT fruits at the time points analyzed (Figure 5). Alanine, aspartic acid, proline, serine, and threonine were notably elevated in *slsamba* fruits, while GABA, glycine, lysine, methionine, phenylalanine, and branched-chain amino acids (valine, leucine, and isoleucine) were reduced in the mutant compared to the WT. It seems likely that these alterations might, at least partially, be related to the rates of protein synthesis essential for cell division.

Concerning organic acids, the levels of fumaric acid, glycolic acid, oxo-glutaric acid, and succinic acid decreased at 8 DPA in *slsamba* mutants, whereas malic acid and quinic acid levels increased compared to the WT. TCA cycle intermediates provide carbon skeletons for the biosynthesis of most amino acids (Galili et al., 2016). Additionally, the accumulation of organic acids during the early stages of fruit development is closely related to the supply of substrates that fuel respiration in this climacteric species (Seymour et al., 2013).

We also identified 47 specialized metabolites via LC-MS (Figure S7) and quantitative enrichment analysis (Figure S7D) revealed that steroidal glycoalkaloids, alcohols and polyols, flavones, flavonoid glycosides, indolyl carboxylic acids and derivatives, flavans and hydroxycinnamic acids and their derivatives were different between *slsamba* and WT fruits. PLS-DA analysis showed that the total variability explained was 77.9% (20.2% from component 1 and 57.7% from component 2) (Figure S7A). Metabolite abundance was visualized with a heatmap (Figure S7B) and the 15 with significant changes (ANOVA P <0.05) are shown in Figure 6. The levels of steroidal glycosides, tryptophan, and caffeoyl glucarate decreased with *slsamba* deletion, while flavone, flavonol, and flavonoid glycosides increased, possibly playing crucial roles in neutralizing reactive oxygen species and modulating responses to pathogens (Frandsen and Narayanasamy, 2018; Chiocchio et al., 2023).

**Figure 6.**
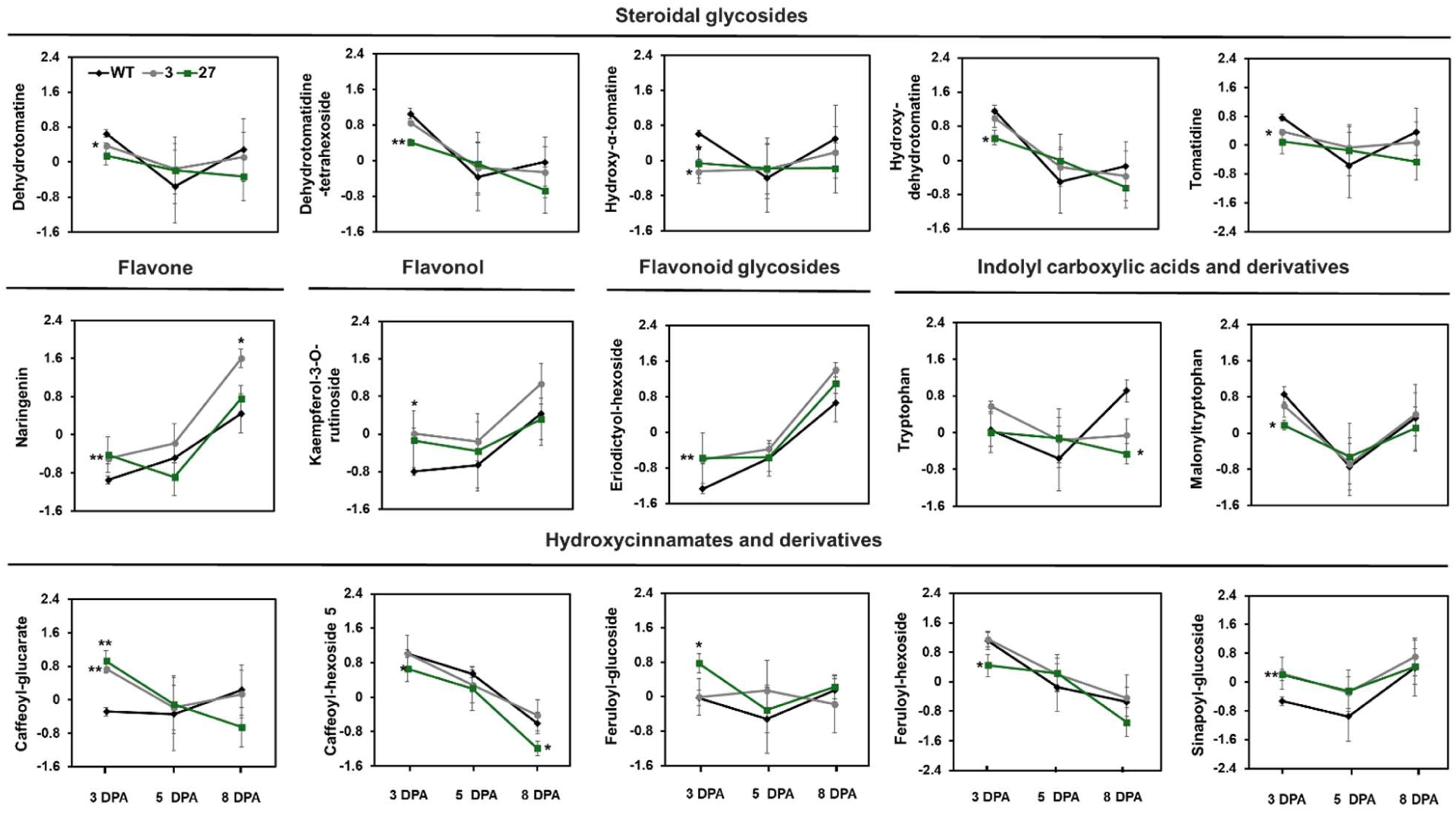
Specialized metabolites significantly altered in fruits of *slsamba* mutants compared to the WT. Relative metabolite quantification was performed by liquid chromatography coupled to tandem mass spectrometry (LC-MS/MS) using fruits from individual homozygous plants of lines 3 and 27 and WT plants harvested at three developmental stages (3-, 5- and 8-days post anthesis). Metabolites were identified based on elution time, molecular weight, and MS/MS fragmentation patterns in our databases (Dataset X) and were normalized using Log and Auto-scaling data on the MetaboAnalyst platform 6.0. Data are means ± SD (n = 4-6). Gray and green lines indicate lines 3 and 27, respectively. Significant differences (one-way ANOVA, post hoc paired t-test) between the WT and each mutant line are indicated by asterisks (*P < 0.05 and **P < 0.01).

### SISAMBA loss of function alters physicochemical parameters of red ripe tomato fruits

To assess whether changes in sugar content observed in the edited *slsamba* lines during early fruit development persisted until fruit ripening, we measured the soluble solids content (°Brix) in red ripe fruits from plants grown in the greenhouse. The °Brix values in *slsamba* were on average 6.3, representing nearly a 20% increase in soluble solids compared to the WT, which had an average of 5.27 (Figure S8). This increase is comparable to the gains achieved by major QTLs associated with improved Brix in tomatoes (Fridman et al., 2004; Zhang et al. 2024).

### Loss of SlSAMBA function leads to cumulative up-regulation of sugar transporter and cell wall modification genes during early tomato fruit development

To examine the effect of *SlSAMBA* disruption at the global transcriptome level, RNA-seq analysis was performed on fruits of *slsamba-3* and WT plants at the same developmental stages as the metabolic profiling. We found 469 differentially expressed genes (DEGs) at 3 DPA, 615 DEGs at 5 DPA, and 4,091 DEGs at 8 DPA (adjusted *P*-value < 0.05 and absolute log_2_(fold change) > 1.5), representing 3.6%, 3.3%, and, 22.8% of the total expressed genes at each time point (Figure S9; complete table in Supplementary Table P1). Several DEGs appeared similarly altered across the three time points, mostly among the down-regulated genes (Figure S9D). At both 5 and 8 DPA the majority of DEGs appeared up-regulated (Figure S9A), and most genes up-regulated at 5 DPA maintained this profile at 8 DPA (Figure S9C). The increase in the number of DEGs over time and the DEGs shared between 5 and 8 DPA suggest that the absence of *SlSAMBA* during the initial stages of fruit development leads to cumulative disturbances in the transcriptional landscape of the tomato fruit.

To shed light on how these transcriptional changes might explain the phenotypes of *slsamba* fruits, we conducted functional enrichment analyses of the DEGs using MapMan4. DEGs were significantly represented in 52 sub-bins belonging to 11 different biological processes, including cell wall organization, protein homeostasis, and solute transport (summarized in Figure 7; complete results in Supplementary Table P2). Similar results were obtained from a functional enrichment analysis of Gene Ontology (GO) terms (Supplementary Table P2).

**Figure 7.**
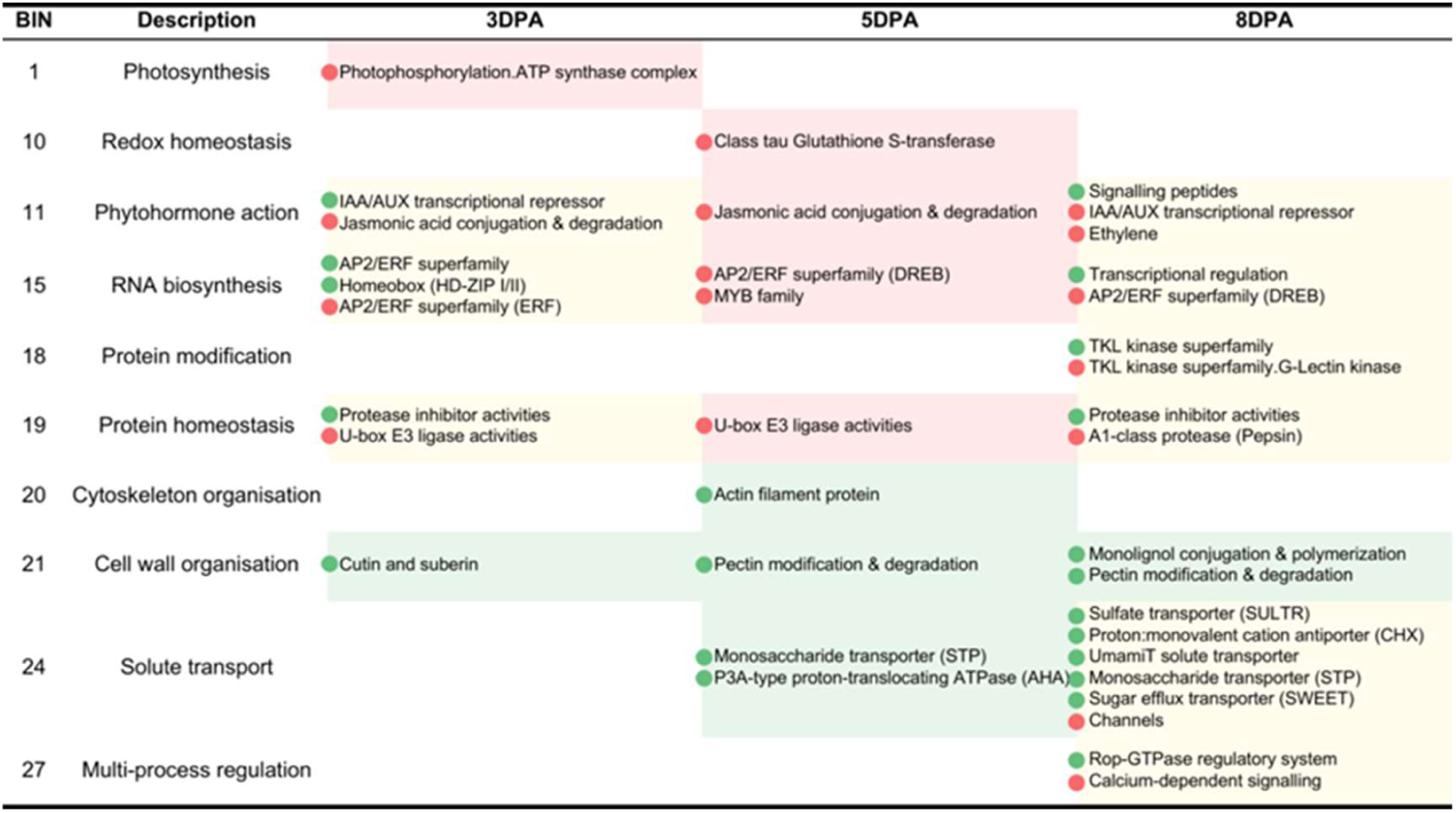
Over-representation analysis of DEGs in MapMan Bins. Enriched top-level MapMan Bins are shown on the left, while specific sub-bins are detailed below each time point. Over-representation was determined using the hypergeometric distribution of DEG lists against the background genes (all genes expressed in the analyzed samples). Only enrichments with Bonferroni-adjusted P-values < 0.05 are portrayed. Up- and down-regulated genes were analyzed separately. Green boxes correspond to an over-represented Bin of up-regulated genes; Red boxes correspond to a Bin enriched in down-regulated genes; Yellow boxes represent Bins over-represented in both up and down-regulated genes. Green circles represent up-regulated bins; Red circles represent down-regulated bins. The complete table with the enrichment analysis results is available in Supplementary Table P1.

Several genes in the solute transport category are involved in sugar transport and were more abundant at 5 and 8 DPA. Upon further examination, we found that eight different sugar transporter gene families were altered in the *slsamba* mutants, totaling 38 DEGs (Supplementary Table P3). This number corresponds to 46.9% of all described sugar transporters in tomato (Reuscher et al., 2014; Feng et al., 2015). Most of these genes were up-regulated at 8 DPA (29 genes, Figure 8), although 15 were already up-regulated at 5 DPA (Figure 8).

**Figure 8.**
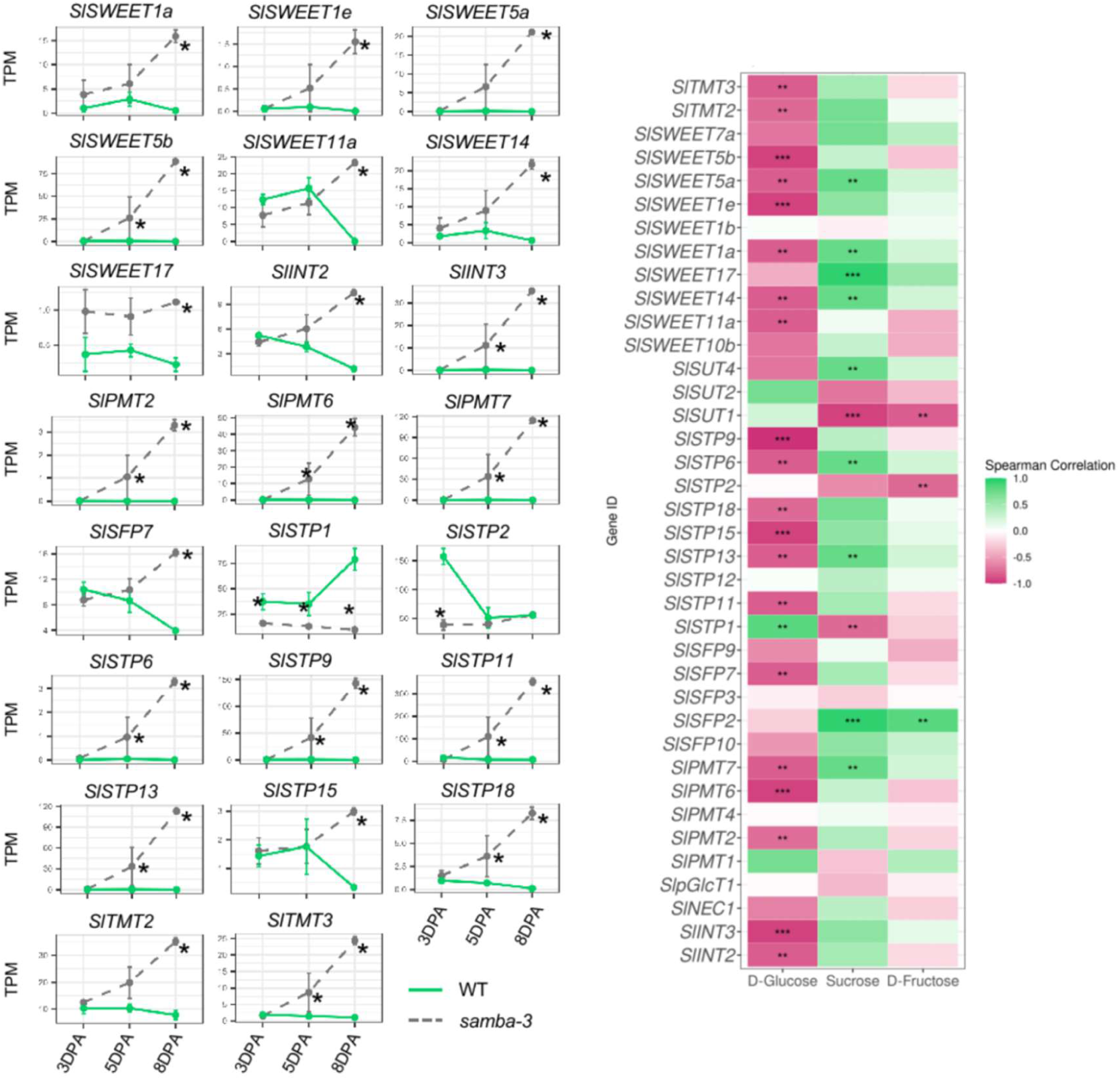
Expression profiles of sugar transporters and their correlation with different sugars. Expression profiles of sugar transporters significantly correlated with different sugars (|⍴| > 0.75). Expression values correspond to normalized TPM values. * = Time point with significant differential expression (adjusted *p-*value < 0.05). The heatmap shows Spearman correlation values (⍴) above 0.75 and below −0.75 marked by ‘**’, while values above 0.9 and below −0.9 are marked by ‘***’.

Similarly, a gene-metabolite correlation analysis with sugar metabolism-related genes revealed an important pattern of expression that may explain the different sugar abundances in *slsamba* fruits (Figure 8). The presence of sucrose metabolism-related genes among the DEGs further consolidates these findings. Notably, at 8 DPA, 13 genes involved in sucrose degradation were up-regulated, including seven beta-fructofuranosidases (invertases), two hexokinases, two fructose kinases, and two sucrose synthases (Supplementary Table P4). Of these, four beta-fructofuranosidases were already up-regulated at 5 DPA: *Solyc09g010090*, *Solyc10g085360*, *Solyc10g085640*, and *Solyc08g079080* (*SlVI2*). These early changes indicate that alterations in sucrose metabolism begin at the initial stages of fruit development, and may contribute to the altered sugar composition observed in the mutants. Our results show that *slsamba-3* fruits exhibit up-regulation of sugar transporters and sucrose degradation enzymes at early developmental stages (5 and 8 DPA), which is unusual for this phase of fruit development (Ruan and Patrick, 1995; Obiadalla-Ali et al., 2004).

A Spearman correlation analysis of the expression of these genes with the abundance of different sugars (Figure 8; Supplementary Table P3) showed that most sugar transporters have a high negative correlation to glucose (⍴ < −0.75) and a high positive correlation with sucrose (⍴ > 0.75). Specifically, 23 out of 38 sugar transporter DEGs showed significant correlations with sugar levels (P < 0.05).

These values indicate that higher expression of these DEGs is associated with lower glucose and higher sucrose content in the fruits, affecting metabolic processes and fruit development.

In the cell wall organization bin, 26 genes were up-regulated at 5 DPA and 138 at 8 DPA, demonstrating a significant and cumulative effect of *SlSAMBA* knockout in cell wall metabolism. Pectin-modifying enzymes were the most enriched in this category at both time points, with 19 up-regulated genes at 5 DPA and 56 at 8 DPA. Since pectin is an intrinsic component of the fruit cell wall and its degradation by pectin-modifying enzymes contributes to fruit softening, these alterations could be linked to the altered fruit shape observed in the mutants. For instance, the up-regulation of genes encoding pectinesterases and polygalacturonases suggests an acceleration of cell wall loosening processes.

Although the cell division Bin did not appear in the MapMan enrichment analysis, several cell cycle-related genes were differentially expressed in the *slsamba-3* fruits, which is expected of an APC/C regulator. The two APC/C activator protein-encoding genes, *CDC20* (*Solyc08g005420*) and *CCS52* (*Solyc12g056490*) were up-regulated at 8 DPA. Moreover, several cyclins were found among the DEGs. At 3 DPA, four cyclins were up-regulated, including *SlCycB1;2* (*Solyc10g080950*), *SlCycB2;4* (*Solyc04g082430*), *SlCycA3;1* (*Solyc12g088530*), and a Cyclin family protein (*Solyc11g030550*). At 8 DPA, four different cyclins were up-regulated: *SlCycA1* (*Solyc11g005090*), *SlCycB2* (*Solyc02g082820*), *SlSDS* (*Solyc04g008070*), and *SlCycU1;1* (*Solyc07g052610*). Additionally, cyclin *SlCycD3* (*Solyc04g078470*) was also up-regulated at both 3 and 8 DPA. The up-regulation of these cyclins, particularly those involved in the G2/M transition, indicates alterations in the proliferation phase of the mutant. These results are strengthened by the cell analysis data, which showed that perturbation of *SlSAMBA* expression also enhanced proliferation in other plant tissues.

As perturbation in the *SlSAMBA* levels leads to alteration in the proliferation status of the plant, genes regulating the activity of the CYC-CDK complex, which controls the cell cycle progression also appeared differentially expressed in *slsamba* fruits. The Kip-related proteins (KRPs) are inhibitory proteins of CDKs, and four genes of this family were altered in our dataset. The *SlKRP5* (*Solyc03g044480*) and *Solyc09g010730* were up-regulated at 8 DPA, while *Solyc09g061280* (*SlKRP2*) and *Solyc09g010980* (*SISMR13*) were down-regulated at this time point. The differential expression of KRPs suggests a complex regulation of CDK activity, potentially balancing the enhanced expression of cyclins. Interestingly, the only gene in this category that appeared at all three time points is *Solyc11g072630*, a mitogen-activated protein kinase (MAPK), which was down-regulated.

To validate the RNA-seq data, we selected 10 genes involved in key metabolic and regulatory pathways, that showed differences in the *slsamba* mutant: tomatidine biosynthesis (*16alpha,22,26-Trihydroxycholesterol*, *Solyc10g018190*), naringenin biosynthesis (*Chalcone synthase SlCHS*, *Solyc01g090600; Chalcone-flavone isomerase SlCHI*, *Solyc05g010310*), sugar metabolism (*Sucrose Synthase SlSUS, Solyc03g098290; Sucrose Phosphate Synthase SlSPS*, *Solyc08g042000*; *Invertase 6 SlINV6*, *Solyc10g0832900*) and transport (*Sugars Will Eventually Be Exported Transporter 5a SWEET5a*, *Solyc03g114200*); and cell cycle regulation (*Cyclin A1 SlCycA1*, *Solyc11g005090*, *Cyclin D3.3 SlCycD3.3*, *Solyc04g078470; Cell Cycle Switch Protein 52B SlCCS52B*, *Solyc12g056490*) (Figure S10). These genes were analyzed using qRT-PCR from *slsamba-3* and WT at −5 and −8 DPA. The qRT-PCR results corroborated the expression patterns observed in the RNA-seq data.

### DEGs in SlSAMBA loss of function plants show enrichment in specific transcription factor (TF)-binding sites

Given the pronounced and cumulative effects of *SlSAMBA* loss of function on gene expression in developing tomato fruit, we investigated whether the *samba* DEGs contain enriched TF binding sites in their regulatory regions. To explore this, we performed a TF-binding site enrichment analysis in the 2-kb upstream sequence of the translation start site of the *slsamba* DEGs, comparing them to all genes expressed at each respective developmental stage sampled (Dataset 2).

Interestingly, within the upregulated DEGs at 3 DPA, our analysis revealed a clear enrichment for motifs of bHLH and homeobox TFs (Table 1). This finding aligns with the GO analysis of the DEGs at 3 DPA, though due to the broad functional roles of bHLH TFs in diverse processes (Hao et al. 2021; Gao et al. 2024), specificity is challenging. Notably, Arabidopsis orthologs of these bHLH and homeobox TFs (specifically from the Glabra subgroup) are known to synergistically regulate metabolic pathways (Nguyen et al., 2023). Conversely, among the downregulated DEGs at 3 DPA, motifs for WRKY and AP2 TFs were enriched, corresponding with the GO analysis of DEGs in which defense categories are enriched (Supplementary Table P2).

**Table 1.** Enrichment of TF-binding sites at timepoints 3 DPA and 5 DPA. Top five TF families with the highest number of enriched motifs, at timepoints 3 DPA and 5 DPA. Expressed TFs are transcription factors known to bind to at least one of the enriched binding sites, which are themselves expressed at the respective timepoint. The **π**-value range indicates the significance of the motif enrichment within each family, where higher values are more significant (combination of adjusted P-value and enrichment fold; see methods).

| TF Family | # Motifs | # Expressed TFs | π-values |
| --- | --- | --- | --- |
| **3 DPA Up-regulated** |  |  |  |
| **bHLH** | 24 | 27 | [1.30-0.74] |
| **Homeodomain** | 5 | 16 | [0.75-0.70] |
| **Sox** | 1 | 7 | [0.31-0.31] |
| **bZIP** | 1 | 6 | [0.75-0.75] |
| **EIN3** | 1 | 5 | [1.10-1.10] |
| **3 DPA Down-regulated** |  |  |  |
| **AP2** | 104 | 40 | [6.29-0.60] |
| **WRKY** | 96 | 31 | [6.30-0.94] |
| **bHLH** | 30 | 29 | [1.92-0.61] |
| **NAC/NAM** | 29 | 13 | [5.42-0.28] |
| **Myb/SANT** | 23 | 17 | [1.78-0.36] |
| **5 DPA Up-regulated** |  |  |  |
| *None* |  |  |  |
| **5 DPA Down-regulated** |  |  |  |
| **WRKY** | 93 | 37 | [11.42-0.54] |
| **AP2** | 88 | 62 | [8.14-0.64] |
| **Sox,WRKY** | 17 | 7 | [4.67-0.84] |
| **NAC/NAM** | 11 | 16 | [5.27-0.56] |
| **E2F** | 10 | 4 | [1.96-0.71] |

At later time points, enrichment is either absent or far less pronounced. Only within the downregulated DEGs at 5 DPA, there is still enrichment for motifs of WRKY and AP2 TFs (Table 1). No enrichment was observed in the upregulated DEGs at 5 DPA, while at 8 DPA (in both up- and downregulated DEGs), different TF classes were enriched (Supplementary Table 5), suggesting that the *slsamba* knockout signal is dispersing and that secondary effects are predominant. This result supports the previous indication that loss of *SlSAMBA* during the early stages of fruit development leads to cumulative disturbances in the transcriptional landscape of the tomato fruit at later stages.

## DISCUSSION

Although the APC/C plays a crucial regulatory role in the eukaryotic cell cycle (Alfieri et al., 2016), knowledge of its function in plants lags behind that of mammals and yeast. Here, we identified *SlSAMBA* as a homolog of *SAMBA* in Arabidopsis, sharing strong similarities with tobacco and exhibiting conservation of key motifs in its protein sequence (Fig. 1). The *SlSAMBA* gene is highly expressed during early flower and fruit development, particularly in floral buds and in fruits at 10 DPA. These stages are associated with intense cell division, supporting its role in cell cycle regulation (Quinet et al., 2019). In the later stages of tomato fruit development, *SlSAMBA* expression decreases and becomes spatially restricted, as observed at anthesis, where it is limited to the stigma and ovary. This pattern differed slightly from that observed in Arabidopsis *SAMBA (AtSAMBA)*, which was highly expressed during embryogenesis, gradually decreasing during seedling germination, and becoming restricted to the hypocotyl after 8 DAS. In later stages of flower development, *AtSAMBA* was also confined to the pollen grains (Eloy et al, 2012). However, unlike in tomato, *SAMBA* was not observed in Arabidopsis female gametophyte, suggesting an additional function for *SlSAMBA* during female gametophyte development specific to the species.

Stable expression of the SlSAMBA-TurboID fusion in tomato roots revealed seven proteins with more than a 4-fold (log2) enrichment, indicating strong proximity or interaction with SlSAMBA. Among the identified proteins, we found homologs of APC6 and APC3b, well-known APC/C subunits, as well as two Arabidopsis homologs of NAP1-RELATED PROTEIN 2 (NRP2), which are also involved in the cell cycle control (Wang et al, 2016). These findings support the hypothesis that SlSAMBA is a member of the APC/C complex, as previously reported for Arabidopsis and maize (Eloy et al., 2012; Gong et al. 2021). The cytoplasmic and nuclear localization of SlSAMBA-eGFP fusion protein observed in transient expression assays supports its regulatory function within the cell. Furthermore, the interaction with APC/C subunits suggests that SlSAMBA may compete with other co-factors, such as CCS52 and CDC20, to modulate substrate selection for APC/C-mediated proteolysis (Kevei et al., 2011; Schmidt et al., 2015).

Using the CRISPR/Cas9 editing system, we generated *slsamba* mutants, which revealed the role of SlSAMBA in coordinating various developmental processes. This was demonstrated by the reduced size of vegetative and reproductive structures, resulting in a dwarf phenotype, as well as altered fruit shape and seed yield. The position of the first inflorescence is lower, resulting in a more compact and dense plant architecture, which has been linked to altered genetic regulation affecting cell proliferation (Xu et al., 2016). *slsamba* mutants have smaller flowers, thinner anthers, and impaired pollen development, which together can impact the reproductive success (Pnueli et al., 1994). Elongated fruit shape is often associated with imbalances in cell proliferation and expansion across the pericarp layers (Xiao et al., 2008; Wu et al., 2018). Disruptions in the APC/C complex in Arabidopsis have been shown to impair gamete viability or embryo formation, ultimately leading to lower seed production (Capron et al, 2003; Kwee and Sundaresan, 2003; Pérez-Pérez et al., 2008; Eloy et al., 2011 Wang et al., 2012, 2013; Guo et al., 2016). This impact on fertility is likely due to impaired cell cycle progression in reproductive structures (Sprunck, 2020; Bolaños-Villegas et al., 2018), a hypothesis supported by studies in other edited tomatoes (Zheng et al., 2022). Compared to WT plants, *slsamba* mutants exhibit reduced overall stature, stem diameter, and leaf area. The latter could directly influence the photosynthetic efficiency of the plant, potentially limiting overall growth and development (Brooks et al., 2014).

Metabolomic analysis of *slsamba* mutants further highlights significant changes in metabolites, particularly amino acids, suggesting a shift in nitrogen distribution potentially regulated by SlSAMBA. This modulation, especially glutamine, a key nitrogen carrier, impacts both fruit growth and flavor characteristics (Beauvoit et al., 2014). In addition, *slsamba* fruits have higher sucrose content and it remains to be elucidated whether this is due to increased fruit sink activity, altered futile cycles involving sucrose synthesis and degradation, or a combination of both. RNA-seq data indicate that the *slsamba-3* mutant shows an early up-regulation of sugar transporter genes and sucrose-degrading enzymes (sucrose synthases, fructokinase, hexokinase, and invertases), suggesting altered sugar transport and accumulation at early developmental stages. Sugars serve as both metabolites and signaling molecules in plant development, potentially influencing the expression of cell cycle genes (Smeekens et al., 2010; Chen et al., 2021). In cultivated tomato varieties, fruits typically accumulate hexoses, with phloem unloading during early development occurring primarily through the symplastic pathway (Ruan and Patrick, 1995; Wang and Ruan, 2013). This symplastic unloading involves the direct transfer of sugars between cells via plasmodesmata, bypassing the need for transporter proteins (Patrick, 1997; Braun, 2022).

As the fruit matures, there is a developmental shift to apoplastic unloading, which requires the activity of invertases to hydrolyze sucrose and hexose transporters to facilitate sugar uptake into cells (Ruan and Patrick, 1995; Wang and Ruan, 2013; Julius et al., 2017). An early shift to apoplastic unloading would typically result in higher hexose levels (Yelle et al., 1988; Dali et al., 1992), which is not the case for *slsamba* fruits. Despite the up-regulation of invertase genes in *slsamba* fruits, the actual enzyme activity might be reduced or inhibited, leading to the observed sugar profile. A recent study on metabolite variation during tomato fruit development (cv Moneymaker) revealed that not always a direct correlation between a metabolite and the transcripts or proteins related to its specific pathway can be done, indicating complex regulations involving coordinated changes at these different levels (Moing et al., 2023).

Sun et al. (2022) reported that differential expression of sugar transporters explains whether a tomato fruit accumulates hexoses or sucrose. By analyzing both sucrose- and hexose-accumulating cherry tomatoes, the authors observed distinct patterns in enzyme activities and transporter expression. In hexose-accumulating tomatoes, high activities of acid invertase, sucrose phosphate synthase (SPS), sucrose synthase (SS), and specific transporters from the SUT and SWEET families are prominent. For sucrose-accumulating fruits, the combination of SPS, SS, and other SUT and SWEET transporters was observed. In our dataset, two *SPS* and two *SS* genes were up-regulated at 5 and 8 DPA, while *SlSUT1* was down-regulated at 3 DPA, *SlSUT2* was down-regulated at 8 DPA, and *SlSUT4* was up-regulated at 8 DPA. Regarding SWEET transporters, the activity of SlSWEET1b, SlSWEET5b, SlSWEET11b, SlSWEET7a, and SlSWEET14 are known to be important for sucrose accumulation (Sun et al., 2022). Of these, only *SlSWEET1b* was not up-regulated in *slsamba* fruits. None of the glucose-accumulating-related SWEET transporters mentioned by Sun et al. (2022) were differentially expressed in *slsamba* fruits. This pattern supports the idea that the up-regulation of specific sugar transporters and metabolic enzymes in *slsamba* fruits contributes to sucrose rather than hexose accumulation in a complex manner.

Elevated intracellular sucrose may serve as a signal influencing gene expression related to the cell cycle (Rawat and Laxmi, 2024), as evidenced by the up-regulation of cyclins such as *SlCycD3* (Solyc04g078470), potentially enhancing cell proliferation activity in early fruit development (Rawat and Laxmi, 2024; Riou-Khamlichi et al., 2000). In Arabidopsis, *CYCD3;1* expression is induced by sucrose (Riou-Khamlichi et al., 2000) and sucrose-starved cells exhibit a decline in CYCD3;1 activity, leading to its subsequent degradation via the proteasome-dependent pathway, resulting in hypophosphorylation of RBR1 and arrest at the G1/S transition (Menges et al., 2006; Hirano et al., 2008; Hirano et al., 2011). In our study, several other cell cycle-related genes, including cyclins involved in the G2/M transition (e.g., SlCycB1;2, SlCycB2;4, SlCycA3;1), were up-regulated in *slsamba-3* fruits. Furthermore, the differential expression of Kip-related proteins (KRPs), which are inhibitors of cyclin-dependent kinases (CDKs), indicates complex regulation of the cell cycle machinery in response to sugar signals (Wang and Ruan, 2013). The up-regulation of some KRPs and the down-regulation of others in *slsamba-3* fruits may represent a balance between promoting and restraining cell division.

Finally, our TF-binding site enrichment analysis suggests that there may indeed exist a specific TF target for SlSAMBA, with two possible scenarios (or a combination thereof). First, SlSAMBA could target an activator-bHLH TF (eventually acting synergistically with a homeobox TF), which in the *slsamba* edited line would be over-accumulating, leading to the up-regulated DEGs at 3 DPA. Second, SlSAMBA would target a repressor TF (which could be either a WRKY and/or an AP2-ERF), which in the *slsamba* edited line would be over-accumulating and leading to the downregulated DEGs at 3 DPA. These findings provide an exciting avenue for future research on the molecular mechanisms by which SlSAMBA modulates fruit development and quality.

In conclusion, our findings demonstrate that SlSAMBA influences sugar transport and metabolism, with changes in intracellular sugar concentrations potentially acting as signals to regulate cell cycle gene expression. This link highlights the dual role of sugars as nutrients and signaling molecules in plant development (Smeekens et al., 2010; Chen et al., 2021). Our findings also suggest the potential for using SAMBA in tomato biotechnology to develop sweeter fruits with higher soluble solids content, a key determinant of taste and economic value (Zhang et al., 2024). Further studies on enzyme activities, sugar signaling pathways, and their impact on cell cycle regulators will provide deeper insights into SlSAMBA’s role in fruit development. Understanding how SlSAMBA disruption leads to coordinated changes in sugar metabolism, cell proliferation, and size could uncover new aspects of fruit growth regulation and the interplay between metabolic and developmental processes.

## MATERIALS AND METHODS

### Sequence alignment and chromosome location

The *S. lycopersicum* SAMBA protein sequence was downloaded from the Genome Database of Solanaceae (http://solgenomics.net/), and the reported SAMBA proteins in *Arabidopsis thaliana*, *Oryza sativa*, *Sorghum bicolor*, *Vitis vinifera*, *Ricinus communis*, *Populus trichocarpa* and *Picea sitchensis* acquired from NCBI. The protein sequence alignment was performed using Clustal X version 2.1 with default parameters (Thompson et al., 1997). The tomato *SAMBA* gene was mapped on a chromosome in accordance with the whole genome of this species. The chromosomal location of the identified *SlSAMBA* gene was extracted from the general feature format (GFF) file provided.

### Plant material and growth conditions

Seeds of *S. lycopersicum* (tomato) cv. Micro-Tom were surface sterilized with commercial bleach (containing an average of 2-2.5% active chlorine) and germinated in half-strength MS medium (Murashige and Skoog; Sigma-Aldrich, St. Louis, MO) supplemented with 30 mg.L^-1^ sucrose and 1% agar, pH 5.8. Seeds were incubated at 25 °C under long-day conditions (16 h light (45 μmol photons.m^-2^.s^-1^ PAR irradiance)/8 h dark). In greenhouse conditions, plants were cultivated with a photoperiod of 16 h/8 h with a mean temperature of 24°C during the day and a mean temperature of 18°C during the night. The humidity was around 55% all day and plants were subjected to daily watering.

### Vector construction and plant transformation

*Solanum lycopersicum* (tomato) cv. Micro-Tom was edited using the CRISPR/Cas9 system to produce *slsamba* mutants. Tomato transformation, CRISPR/Cas9 vector, and primers are described in Supporting Information Table X. T0 generations of the mutants were backcrossed to wild-type (WT) once or twice to separate alleles and select against Cas9. Mutations were confirmed by Sanger sequencing. The Cas9-free homozygous mutants were obtained in the F2 or F3 generations and used for further analysis.

For the *SlSAMBA* promoter analysis, a 1,600 bp genomic fragment (upstream of the ATG start codon) containing the putative promoter region was amplified from the genomic DNA of Micro-Tom tomato plants, cloned into the pDONR221 vector (Invitrogen), and subcloned into the GUS::GFP-containing binary vector pKGWFS7. Micro-Tom transgenic plants were transformed using *Agrobacterium tumefaciens* strain GV3101 as described by Pino et al. (2010). Five independent antibiotic-resistant transgenic lines were selected by PCR and were subsequently examined for expression levels.

### Subcellular localization of the SlSAMBA protein

Transient expression in *N. benthamiana* leaves via agroinfiltration was performed as described by Zhang et al. (2020). *Agrobacterium tumefaciens* strain harboring the N-terminal GFP fusion vector was grown in LB medium in the presence of antibiotics. The agrobacteria culture was pelleted and resuspended in the infiltration medium (¼MS pH 6.0, 1% Sucrose, 100 μM Acetosyringone, 0.005% (v/v) Silwet L-77) adjusted to an OD_600_ of 0.5. Agrobacteria suspension was infiltrated into the abaxial side of the tobacco leaves using a needless syringe. Images were captured by confocal laser scanning microscopy (Leica TCSSP5 AOBS, Wetzla, Germany). Excitation and emission wavelengths for GFP were 488 nm and 505–530 nm, respectively. The subcellular localization was predicted by using the online tools CELLO v.2.5 (http://cello.life.nctu.edu.tw/) and WoLF PSORT (https://wolfpsort.hgc.jp/). Transmembrane prediction was carried out with HMMTOP software.

### RNA extraction and gene expression analysis

Total RNA was isolated from leaf tissues of tomato cv. Micro-Tom using TRIzol (ThermoFisher, USA), according to the manufacturer’s instructions. RNA was quantified using NanoDrop 2000 spectrophotometer (Thermo Scientific, USA), and its integrity was checked by electrophoresis. To eliminate the residual genomic DNA present in the preparation, the RNA was treated by RNAse-free DNase I according to the manufacturer’s instructions (Promega, USA).

The cDNA was synthesized from 1 μg of total RNA using the SuperScript™ III First-Strand Synthesis System, according to the manufacturer’s protocol (ThermoFisher, USA). Transgene expression was analyzed by qRT-PCR on an ABI 7500 qPCR thermocycler (Applied Biosystems, USA) using the Platinum SYBR Green Supermix (Invitrogen, USA). Three biological replicates and three technical replicates for each reaction were analyzed. The reaction was initiated at 95 °C for 10 min, and followed by 40 amplification cycles of 95°C for 15 s, 60°C for 30 s, and 72°C for 30 s. The target was the *SISAMBA* (Table S4) and *SIβ-ACTIN* was used as a reference gene, following previous works (Ferreira Silva et al. 2014) (Table S4). The comparative ΔΔCt method was used to calculate relative expression levels in Real-Time qPCR data (Livak and Schmittgen 2001). Changes in gene expression related to sugar synthesis, flavone synthesis, and cell cycle were also characterized by qRT-PCR using the primers listed in Table S4

### Histochemical GUS assay

Flowers at anthesis and fruits at 10, 15, and 30 DPA and red ripe (RR) fruit from pSlSAMBA::GUS::GFP lines were harvested. Samples were incubated in 50 mM sodium phosphate buffer (pH 7.0) containing 0.9 mM 5-bromo-4-chloro-3-indolyl-beta-D-glucuronide (X-Gluc), 10 mM EDTA, and 0.1% (v/v) Triton X-100 at 37°C for 2 h for GUS staining. Subsequently, the stained samples were treated with 70% ethanol at 37°C for 2 days to remove chlorophyll. Images were acquired using the Axio Zoom V16 fluorescence microscope (Zeiss, Germany).

### Turbo ID-catalyzed proximity labeling

TurboID-mediated proximity labeling involved transformed tomato hairy root cultivation with *Agrobacterium rhizogenes*, sample preparation for mass spectrometry (MS), immunoblot analysis, LC-MS/MS analysis, and MS-data analysis, which were all carried out essentially as described (Gryffroy et al., 2023a and 2023b).

### Yeast two-hybrid assays

The full-length coding sequences (CDS) of *SlAPC10* (*Solyc02g062680*), *SlSAMBA* (*Solyc08g076580.2*), and *SlCDC27B* (*Solyc03g0203431*) were cloned into the pGADT7 and pGBT9 Gateway™ vectors (Cuéllar et al., 2013). All CDCS were cloned in both vectors. The pGADT7 vector contains the activation domain (AD) of the GAL4 transcriptional activator, while pGBT9 includes the GAL4 DNA-binding domain (BD). The destination vectors were co-transformed into the *Saccharomyces cerevisiae* PJ69-4A strain, using Frozen-EZ Yeast Transformation II Kit (Zymo Research, California, USA). Transformants were selected on SD medium (Takara Bio, Shigo, Japan) lacking leucine and tryptophan (SD -Leu -Trp). Three individual colonies were chosen, grown overnight in liquid culture at 30°C, under 250 rpm agitation, and sequentially diluted (10- to 100-fold). The dilutions were dropped on both control (SD -Leu -Trp) and selective media lacking leucine, tryptophan, and histidine (SD -Leu -Trp - His). Plates were incubated at 30° C for 3 days.

### Phenotypic analyses

To evaluate morphological diversity among the *slsamba* and wild-type (WT) plants, the height of the first inflorescence and stem diameter were measured in 40-day-old CRISPR-Cas mutants (T2) and WT plants (12 biological replicates (one plant each) per genotype). The morphology of the 5^th^ leaf (21 biological replicates (single leaves) per genotype) and total leaflet area were measured 1 month after sowing. Flowers at anthesis (21 biological replicates (flowers) per genotype) were observed by stereo microscopy (SMZ 1500 increased 7.5×). The length and width area were quantified using ImageJ (https://imagej.nih.gov/ij/).

### Pollen quantification and viability

Pollen fertility was determined by counting the number of pollen grains stained in red with acetic carmine (2%) solution (Kearns and Nouye, 1993). Anthers from the flower buds were dissected and pollens were extracted with 500 mL of 0.5 M sucrose, pelleted, resuspended in 200 mL, deposited on a microscope slide, and stained. The number of pollen grains was counted in 10 randomly selected fields per slide. The percentage of pollen fertility was obtained using (number of viable pollen grains)/(number of total pollen grains counted × 100)].

Pollen germination tests were performed on glass slides coated with germination medium [0.292 M sucrose, 1.27 mM Ca(NO_3_)_2_, 1.62 mM H_3_BO_3_,1 mM KH_2_PO_4_, and 0.5% agarose]. The number of germinated pollen grains was counted with a microscope after 2 h of incubation at 25°C in the dark. Pollens were observed using a bright field microscope (Zeiss, Axioplan) and photographed with a CCD camera (Motic 3 megapixels). The percentage of pollen germination was calculated by the number of germinated pollen grains/number of total pollen grains counted × 100.

### Scanning transmission electron microscopy (STEM)

Pollen grains from *slsamba* and WT plants were prepared for STEM. After air-drying, the pollen grains were mounted on a sample plate and coated with a 20 nm-thick layer of gold using a Hummer VII-Sputter facility (Analtech). Subsequently, the samples were examined under a ZEISS Gemini SEM 300 scanning electron microscope operating at 30 kV. Image acquisition was performed using a Mamiya RB67 camera connected to the microscope.

### Fruit growth parameters and cytological analysis

Fruits from 12 plants per genotype were harvested at anthesis, at 10-, 15-, 20-, 25-, and 30 DPA, and at the RR stage. Fruits were weighed on a semi-analytical scale and imaged with a Nikon D5300 camera before being cut at the equatorial region. The pictures were analyzed with ImageJ (https://imagej.nih.gov/ij/) software to measure fruit diameters, fruit height, and pericarp thickness. Fruit diameters were measured at the equatorial region (D_eq_) in two directions and averaged. The shape index was calculated as the ratio of fruit height and fruit D_eq_. The pericarp thickness corresponds to the average of six measurements distributed around the equatorial section. Per genotype, 24 biological replicates (fruits) were analyzed. Equatorial pericarp fragments were fixed in FAA (formaldehyde 4%, ethanol 50%, acetic acid 5%) by applying a strong vacuum of 400 mmHg for 15 min. The fixative was renewed, and the samples were incubated overnight at 4°C. Pericarps were sliced at 150 μm thickness using an HM 650V Vibrating-Blade Microtome (Thermo Scientific). The sections were transferred to phosphate buffer saline (PBS, consisting of 137 mM NaCl, 2.7 mM KCl, 10 mM Na_2_HPO_4_, 1.8 mM KH_2_PO_4_, pH 7). PBS was replaced by a PBS-staining solution containing 4’,6-diamidino-2-phenylindole (DAPI) at 20 μg/mL and calcofluor white at 1.33 μg/mL following incubation for 15 min. After three washes of 5 min with PBS, the sections were mounted with Citifluor™ AF1 (Electron Microscopy Science) to reduce fluorescence fading. Sections were imaged with a Zeiss LSM880 confocal laser scanning microscope using a 20x dry objective (NA 0.8). For calcofluor and DAPI visualization, excitation was performed at 405 nm and fluorescence emission was collected at 420-480nm. Cell walls were manually outlined using SketchBook software and cell size was determined using Image J. The number of cell layers was counted.

### Ovary growth parameters and cytological analysis

Flowers at anthesis and stage 11 (4 mm) (Brukhin et al., 2003) from 24 plants per genotype were harvested. The perianth and anthers were removed keeping the ovary fixed on the peduncle to facilitate manipulation. Pictures were taken to measure diameter and length, and then samples were immediately immersed in FAA by applying a vacuum of 350 mmHg for 15 min. The fixative solution was renewed, and samples were incubated overnight at 4°C. Samples were dehydrated using an ethanol series (70, 96, 100%), transferred to histosol, and finally embedded in paraffin wax. Ovaries were sectioned (8 μm thick) in longitudinal and transversal directions and stained with 0.1% toluidine blue. Slices were imaged with an Axiozoom macroscope (Zeiss) and analyzed with ImageJ software. Three regions of each ovary wall were analyzed for measuring thickness, counting the number of cell layers, and measuring the cell size. The average cell size was determined by the ratio of a selected area to the number of cells.

### Ploidy level analysis

Frozen equatorial sections of fruit pericarps (*slsamba* and WT) of 0, 3, 5, 10, 15, 20, 25, 30 DPA, and red ripe were chopped with a razor blade into 400 μl of chilled CyStain UV Precise P Nuclei Extraction Buffer (Sysmex). The suspension was filtered through a 50 μm nylon filter, and 1,600 μL of chilled CyStain UV Precise P Staining Buffer (Sysmex) was added to the isolated nuclei. The nuclei DNA content was measured using a 208 CyFlow 9 Space flow cytometer (Sysmex). Ploidy profiles were then analyzed with FloMax software (Sysmex). The percentage of all ploidy levels was calculated from the raw count obtained through the gating region of each peak. The 2C nuclei in the 30 DPA and red ripe stages were set to zero as the corresponding peaks are in the background noise. The mean C value (MCV) defined as the sum of each C value class weighed by their frequency was calculated (Cheniclet et al., 2005).

### Fruit quality

Red ripe fruits (52 DPA, 24 fruits per genotype) were homogenized and the total soluble solids (°Brix) of the resulting juices were measured with a digital Brix refractometer (ATAGO PAL-BX/ACID3).

### Extraction and analysis of metabolites

Metabolite analyses were performed in fruits from WT and individual homozygous *slsamba* plants (lines 3 and 27) harvested at three developmental stages (3-, 5-, and 8 DPA). Pools of 10 fruits were collected, immediately frozen in liquid nitrogen, powdered, and stored at −80°C until extraction.

Extraction and quantification of primary and secondary metabolites (4-6 replicates) were performed as described by Salem et al. (2016). Briefly, 150 and 300 µL of the polar phase were dried in a centrifugal vacuum concentrator for primary and secondary metabolite profiling, respectively. The primary metabolite pellet was resuspended in 40 µL methoxyaminhydrochloride (20 mg.mL^-1^ in pyridine) and derivatized for 2 h at 37°C. Afterward, 70 µL of N-methyl-N-[trimethylsilyl] trifluoroacetamide containing 20 µL.mL^-1^ fatty acid methyl esters mixture as retention time standards were added. The mixture was incubated for 30 min at 37°C at 400 rpm shaking. A volume of 1 µL of this solution was injected into an Agilent 6890N gas chromatograph coupled with a LECO Pegasus III time of flight mass spectrometry (TOF-MS) running in electron ionization (EI) + mode.

The secondary metabolite pellet was resuspended in 200 µL 50% (v/v) methanol in water and 2 mL was injected into RP high-strength silica T3 C18 column using a Waters Acquity UPLC system. The analysis workflow included peak detection, retention time alignment, and removal of chemical noise. PLS-DA and heat map analysis were performed by MetaboAnalyst 6.0 (https://www.metaboanalyst.ca/).

### RNA-seq

RNA-seq analysis was performed on three pools of the same samples analyzed for primary and secondary metabolites. Total RNA was isolated using TRIzol (ThermoFisher, USA) according to the manufacturer’s instructions, purified and quantified with a NanoDrop 2000 spectrophotometer (Thermo Scientific, USA) and Qubit, and its integrity was examined by electrophoresis. To eliminate the residual genomic DNA present in the preparation, the RNA was treated by RNAse-free DNase I according to the manufacturer’s instructions (Promega, USA).

RNA samples were sent to the NGS services Fasteris Co., Ltd. (Switzerland) for sequencing. The sequencing libraries were prepared with the Illumina TruSeq stranded mRNA kit, and sequenced in 150 bp paired-end mode in NovaSeq 6000. At least 50 million read pairs were obtained for each sample. Each sequencing library was then processed with *fastp v.0.23.4* (Chen et al., 2018) to remove adapters, poly-A tails, reads with more than 20% bases below Phred quality of 30, and reads smaller than 100 bp and larger than 150 bp. The libraries’ strandedness was checked with the software *how_are_we_stranded_here v.1.0.1* (https://github.com/signalbash/how_are_we_stranded_here) before quantification.

Transcript quantification was performed with *salmon v.1.10.0* (Patro et al., 2017) using the gentrome as a reference, a combination of the transcripts (ITAG4.1) and genomic (SL4.0) sequences where the latter are used as decoys to avoid mismapping. From this point forward, the analyses were performed individually for each time point. The quantification files were imported into R version 4.4.1 with the package *tximport 1.32.0* (Soneson et al., 2015) to generate transcript- and gene-level count tables. Since ITAG4.1 only provides a single transcript per gene, all analyses were performed at the gene level. Genes with low expression were filtered from the count table with a cutoff of 1 CPM in at least three samples. Then, the filtered counts were normalized by TMM (Robinson and Oshlack, 2010), and the mean-variance relationships for each gene were calculated with voom using sample weights to account for sample heterogeneity (Liu et al., 2015). The limma package was used to fit a linear model, apply contrasts between the genotypes, and adjust variance estimates (Ritchie et al., 2015). The results of the DE analysis were filtered by choosing an adjusted P-value < 0.05 and an absolute log2(fold-change) of 1.5.

For metabolite-gene correlation analyses, the average normalized expression values of genes and metabolites at each time point were used. Then, Spearman’s correlation coefficient was calculated using R version 4.4.1. Correlation values (⍴) above 0.75 and below −0.75 were considered relevant for the interpretation of the results.

### TF-Binding Site Enrichment Analysis

Enrichment analysis for TF-binding sites (TFBS; or motifs), in 2-kb sequences upstream from the translation start site of the *slsamba* DEGs, was performed essentially as described by Gryffroy et al. (2023a). Where needed, the upstream sequences were shortened to avoid overlap with neighboring genes. Known motifs for tomato were collected from CisBP version 2.00 (Weirauch et al., 2014) and JASPAR 2020 (Fornes et al., 2020). These motifs were mapped onto the 2-kb sequences upstream of *slsamba* DEGs using FIMO (MEME version 4.11.4; default parameters) (Grant et al., 2011) and Cluster-Buster (compiled on September 22, 2017) (Frith et al., 2003) with cluster score threshold (−c) set to 0. Following Kulkarni et al. (2024), the 7,000 top-scoring motif matches from FIMO were combined with the 4,000 top-scoring matches from Cluster-Buster, and lower-scoring matches were discarded. Within each set of DEGs, a hypergeometric test was performed per motif, and all motifs with Benjamini-Hochberg adjusted P-value < 0.05 were considered enriched within that DEG set. To focus on motifs that are specifically enriched in the set of up- or down-regulated genes at a certain time point, the set of expressed genes (with known motifs in their 2-kb upstream sequence) at that time point was used as background for the hypergeometric tests. To sort enriched motifs, taking into account both statistical significance (adjusted P-value) and enrichment fold, π-values were computed following Xiao et al. (2014). Enriched motifs were sorted decreasingly by π-values.

### Statistical analysis

All values were expressed as the mean ± standard Error of the Mean (SEM). Data from lines *slsamba* and WT plants were analyzed with ANOVA and Dunnett’s tests using GraphPad Prism 5 software (La Jolla, CA). Probabilities of P < 0.05 and P < 0.01 were considered statistically significant.

## SUPPLEMENTARY MATERIAL

**Figure S1.**
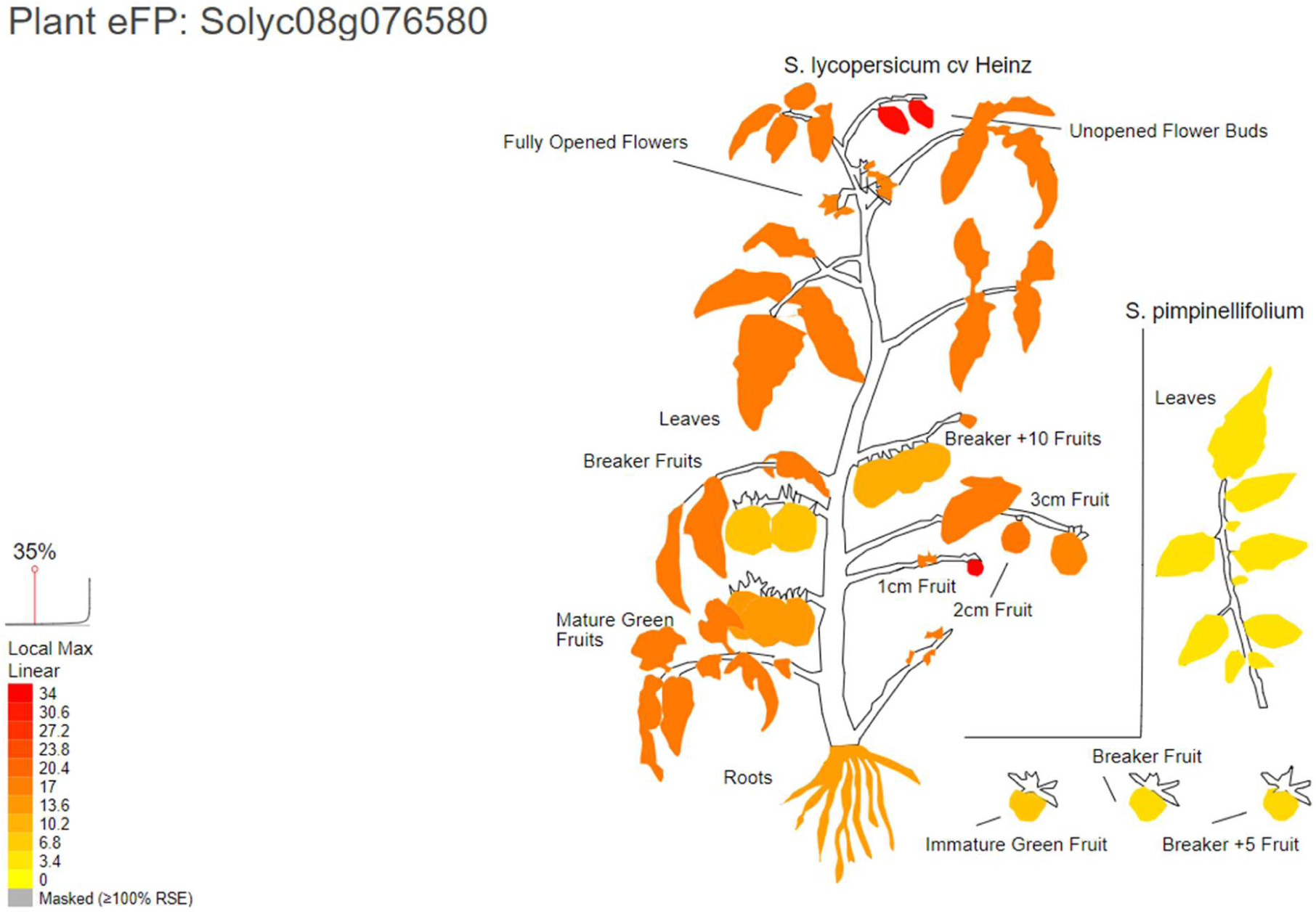
Plant BAR eFP browser showing the expression of *SAMBA* (*Solyc08g076580*) in distinct tomato tissues. Image extracted from the BAR ePlant tomato browser (https://bar.utoronto.ca/eplant_tomato/).

**Figure S2.**
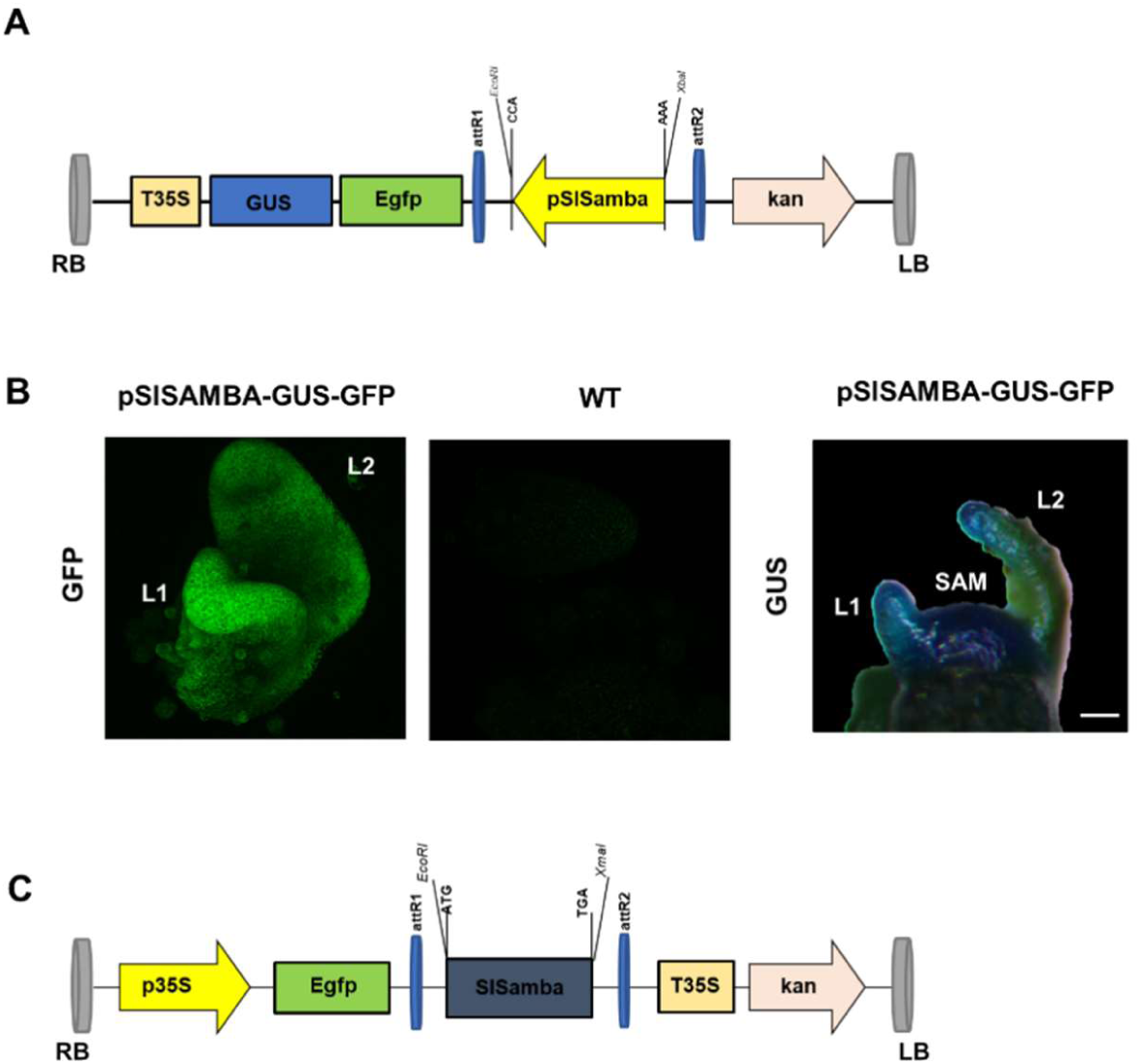
Expression of pSISAMBA-GUS-GFP and SAMBA:EGFP vectors in tobacco. (A) To study the dynamic expression of *SlSAMBA*, a 1.6-kb fragment upstream of the ATG codon of the *SISAMBA* gene was inserted in frame into EcoRI and Xmal sites of pKGWFS7 upstream EGFP and GUS coding region and introduced into tomato plants. The pKGWFS7 contains the selective marker *neomycin phosphotransferase II* (*nptII*) that gives resistance to kanamycin. (B) *SlSAMBA*-promoter-directed expression of GFP and GUS in the early vegetative meristem (7-day-old) of tomato transformed plants (T3). Images of tomato apices showing vegetative meristem observed under fluorescence excitation (GFP) with a confocal microscope. Leaf primordium (L) and shoot apical meristem (SAM). The right lower panel shows the WT vegetative meristem without almost fluorescence. *SAMBA*-promoter-directed expression of GUS in tomato transformed plants (T3). (C) Vector used in the subcellular localization of SAMBA:EGFP (N-terminal) fusion in epidermal cells of *Nicotiana benthamiana*.

**Figure S3.**
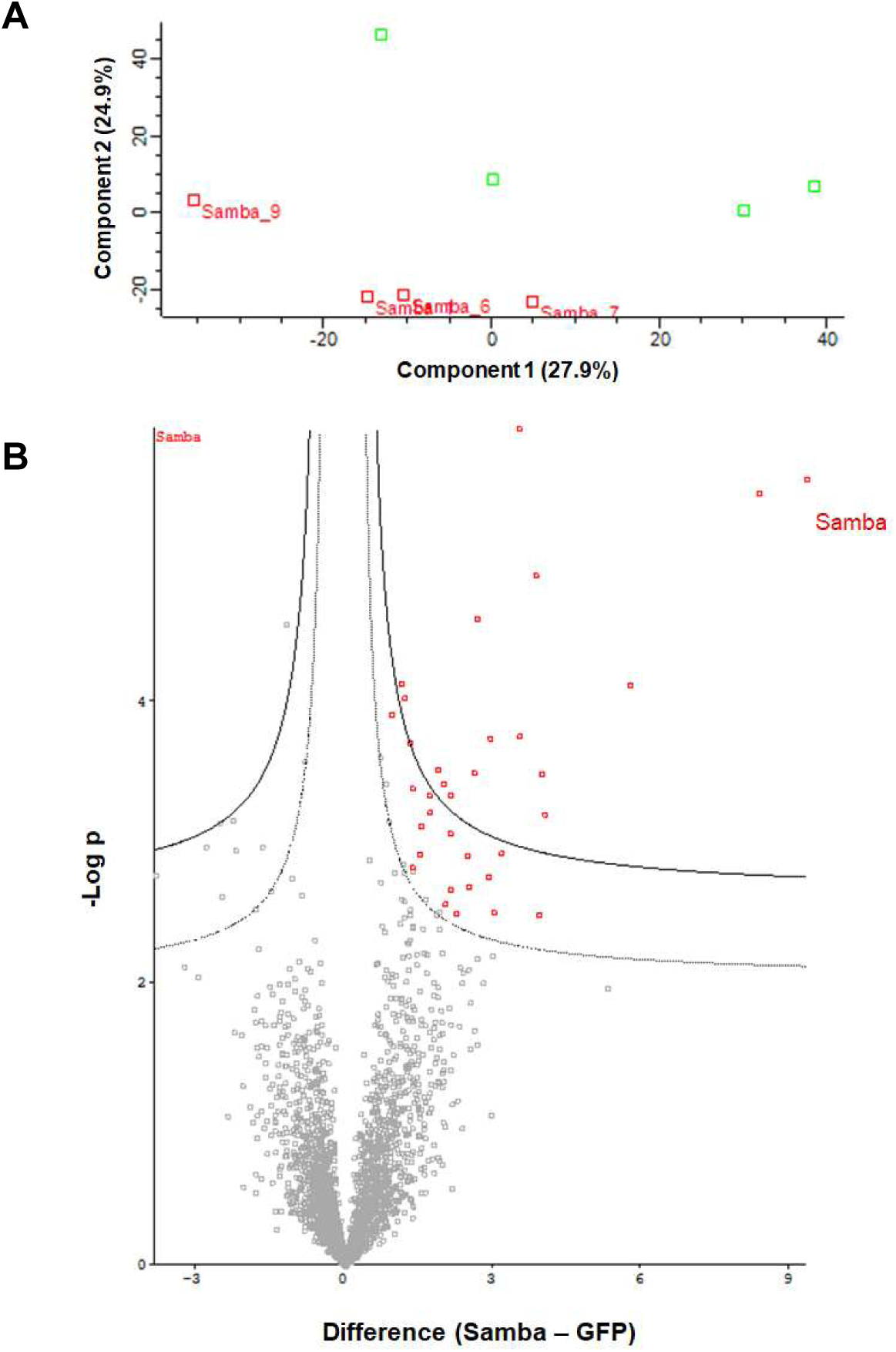
Statistical analysis on SlSAMBA, eGFP TurboID samples, and SlSAMBA interacts with CDC27b in the yeast two-hybrid (Y2H) assay. *35S::GFP-TurboID* and *XVE::samba-TurboID* expression constructs were used for the rhizogenic *Agrobacterium*-mediated transformation of tomato. Transformed hairy roots were treated with β-estradiol (100 μM) for 24 h and biotin (50 μM) for 2 h. Proteins were extracted from the hairy root tissue, enriched with streptavidin beads, digested with trypsin, and identified by mass spectrometry. The MaxQuant software was used for peptide and protein identification on the acquired raw files and the Perseus software for statistical data analysis (Cox and Mann, 2008; Tyanova et al., 2016). (A) Sample variability represented by a principal component analysis (PCA) plot. Red circles and green squares are SAMBA and eGFP samples, respectively. (B) Volcano plot of pairwise comparison between SAMBA and eGFP samples. A two-sample Student’s t test was done to identify enriched proteins in the Samba samples. The full line indicates the cut-off at false discovery rate (FDR)=0.01 and S0=0.1 (i.e., artificial within-group variance; which defines the relative importance of the P value and difference between means). The t test difference was plotted against the t test –log (*P* value). (C) **Y2H**

**Figure S4.**
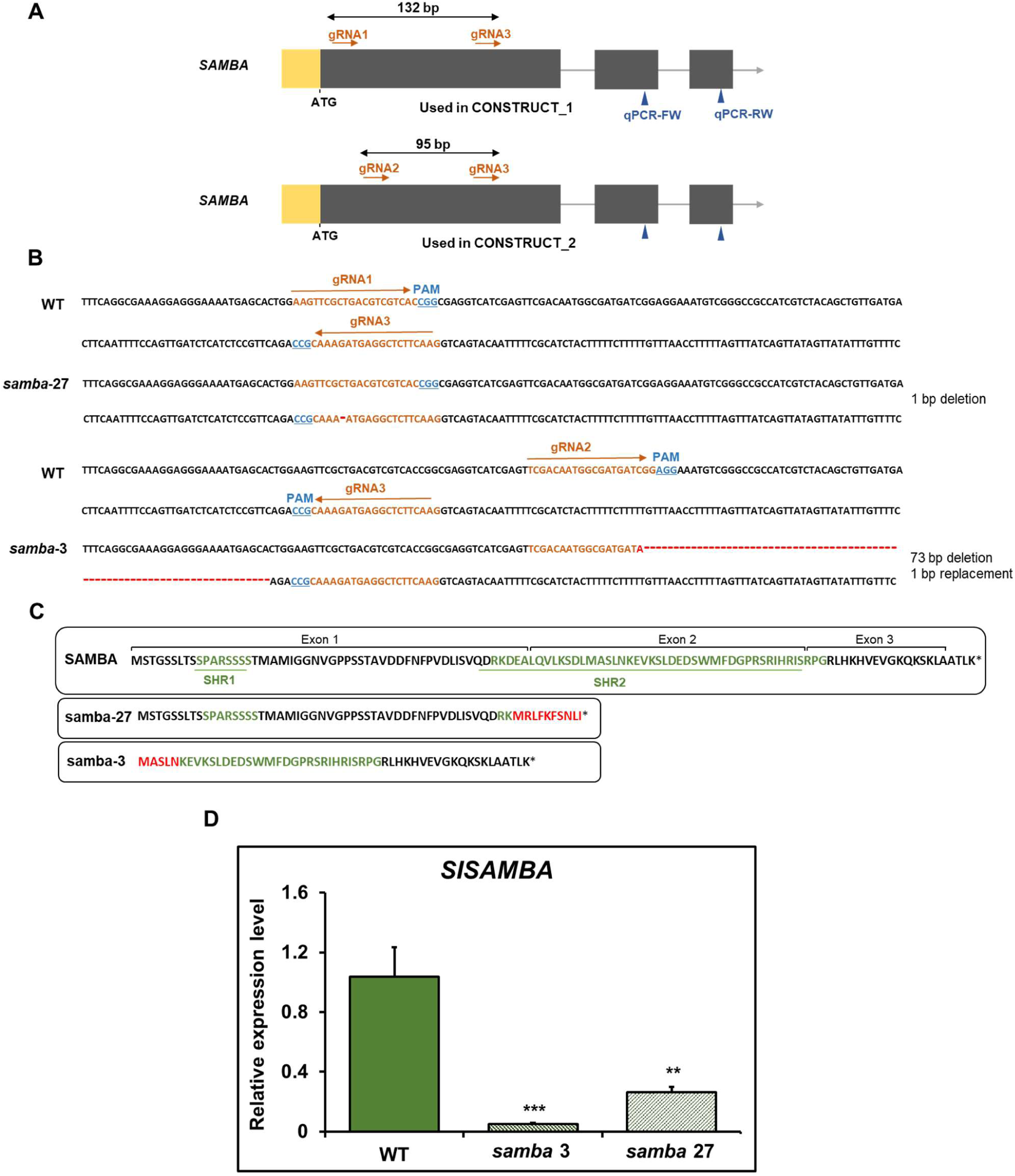
Generation of *samba* mutants by CRISPR/Cas9. (A) Structural representation of the *SISAMBA* gene used in this study, showing the different target sites of the two gRNAs used per construct. UTR, exons, and introns are indicated by yellow, black boxes, and grey bars, respectively. qPCR-FW and qPCR-RW indicate the position of the forward and reverse primers used for *SISAMBA* quantitative RT-PCR (qRT-PCR). (B) The wild-type (WT) sequence is shown with the gRNA sequences highlighted in orange and the PAM sequence in blue. #3 and #27 are the mutants obtained in this study. The mutation sites are shown in red. (C) Amino acid sequences of SISAMBA in WT and *samba* mutant isoforms. SAMBA homology region 1 (SHR1) and SHR2, as defined by Eloy et al. (2012), are marked in green and the missense amino acids are indicated in red. Asterisks represent a stop codon. (D) qRT-PCR transcript analysis of wild-type (WT) and *slsamba* plants (#3 and #27). Total RNA was prepared from the whole seedlings harvested 30 days after sowing (DAS) and amplified by qRT-PCR. All values were normalized against the expression level of the *β-ACTIN*. Data are means ± SD (n = 3). Significant differences (ANOVA followed by Dunnett’s test) are indicated by asterisks (*P < 0.05 and **P < 0.01).

**Figure S5.**
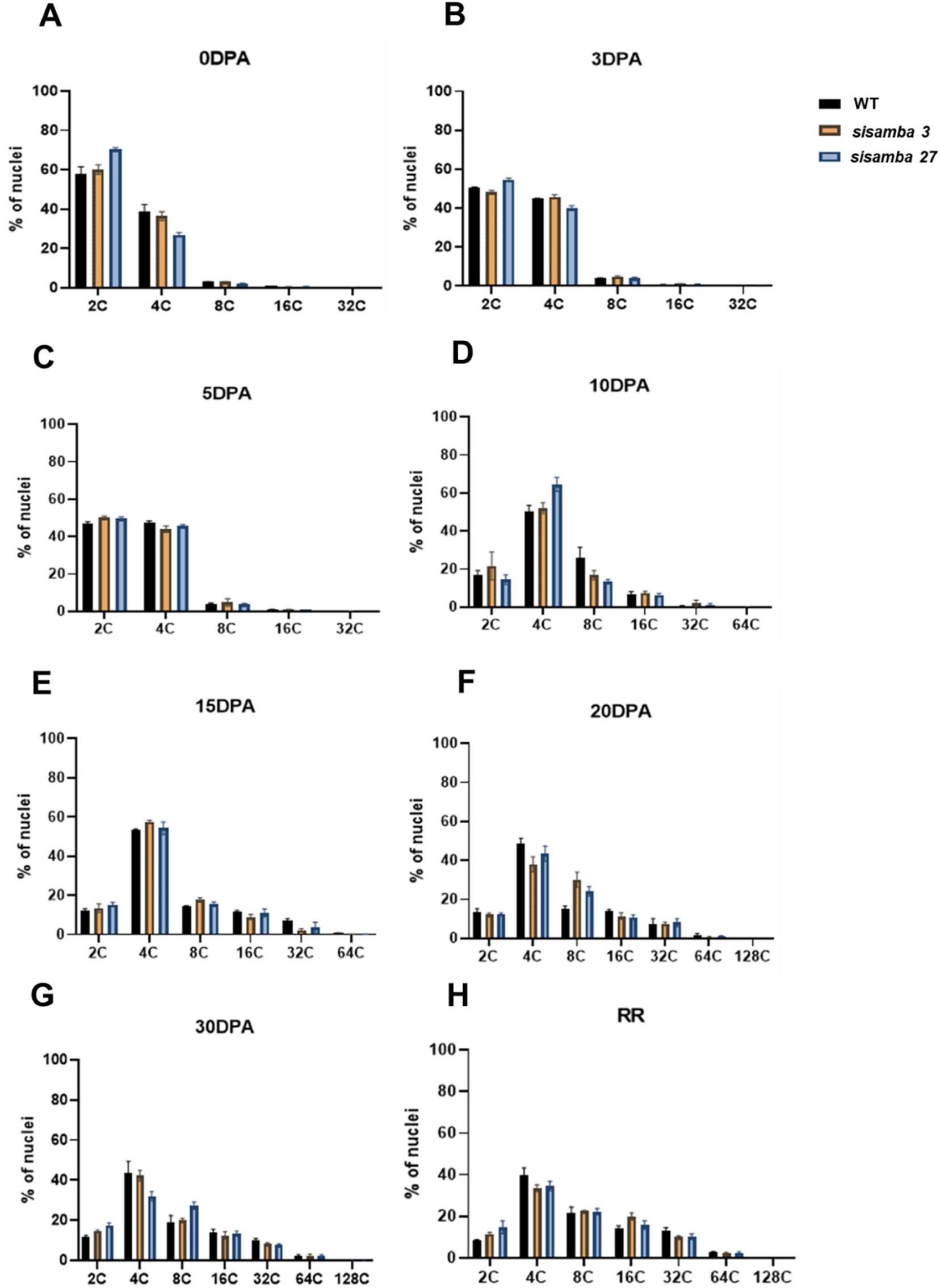
Flow cytometric analysis of nuclear DNA ploidy distribution in pericarps of green and red ripe wild-type (WT) and *slsamba* fruits (#3 and #27). Nuclei were isolated from pericarps of 0, 3, 5, 10, 15, 20, and 30 days after pollination (DPA) green (A -G) and 58 DPA red ripe tomato fruit (H) using chilled CyStain UV Precise P Nuclei Extraction buffer. Nuclei were stained by DAPI (4, 6-diamidino-2-phenylindole) at the final concentration of 50 µg/ml for flow cytometric analysis.

**Figure S6.**
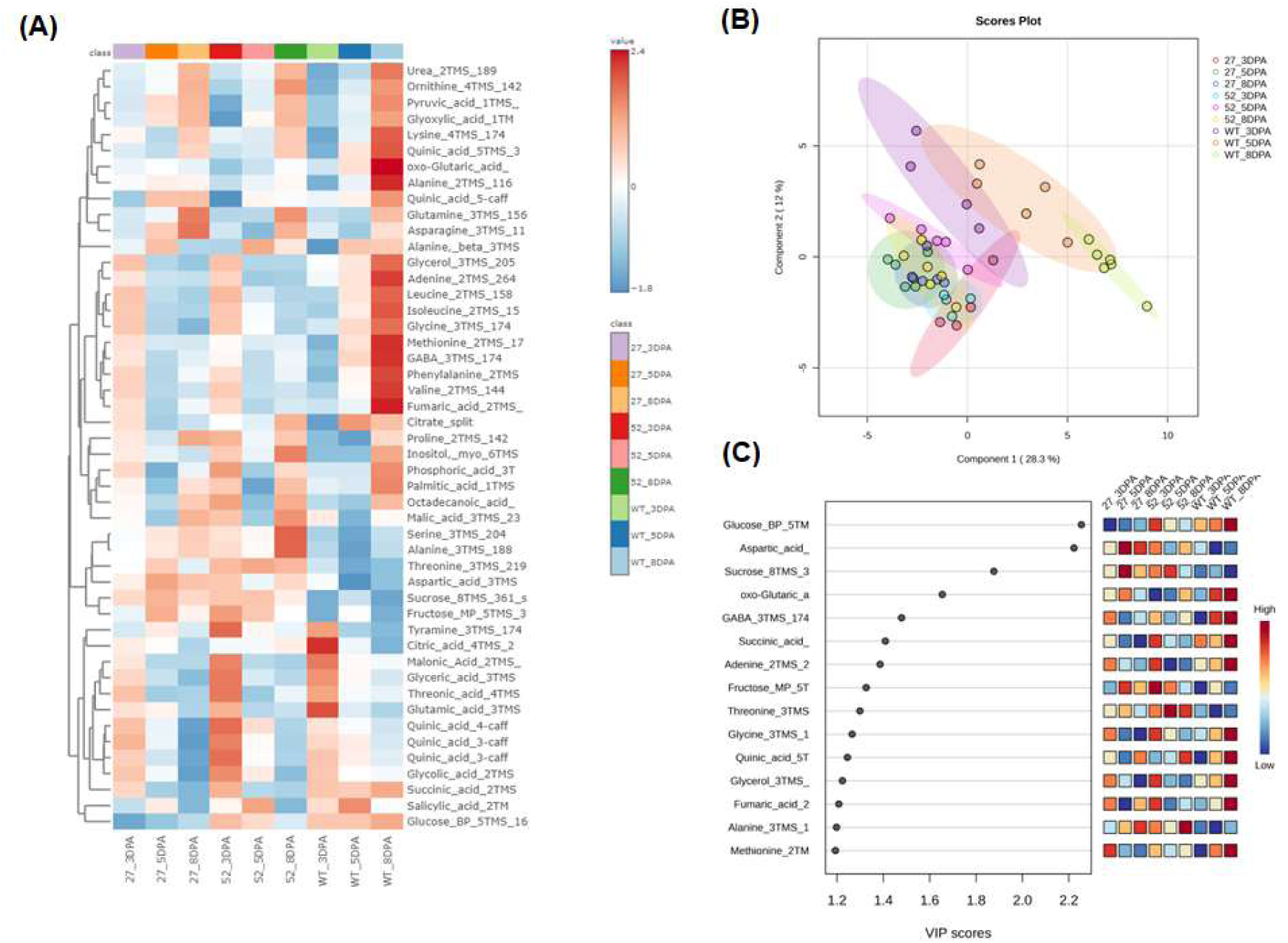
Metabolic characterization of tomato fruit at three developmental stages −3, −5, and 8 days post anthesis (DPA) collected and analyzed by GC–MS. (A) Heat map of the metabolites identified in this study. (B) Partial least square-discriminant analysis (PLS-DA). (C) Variable importance in projection (VIP) scores of the PLS-DA model. Metabolites included in this VIP score list have scores higher than 1, which indicates those that mostly contributed to the separation observed in the PLS-DA model. PLS-DA was carried out by combining data from all stages of development. The data were normalized by using Log and Auto-scaling transformations on the MetaboAnalyst platform 6.0 (n = 4-6).

**Figure S7.**
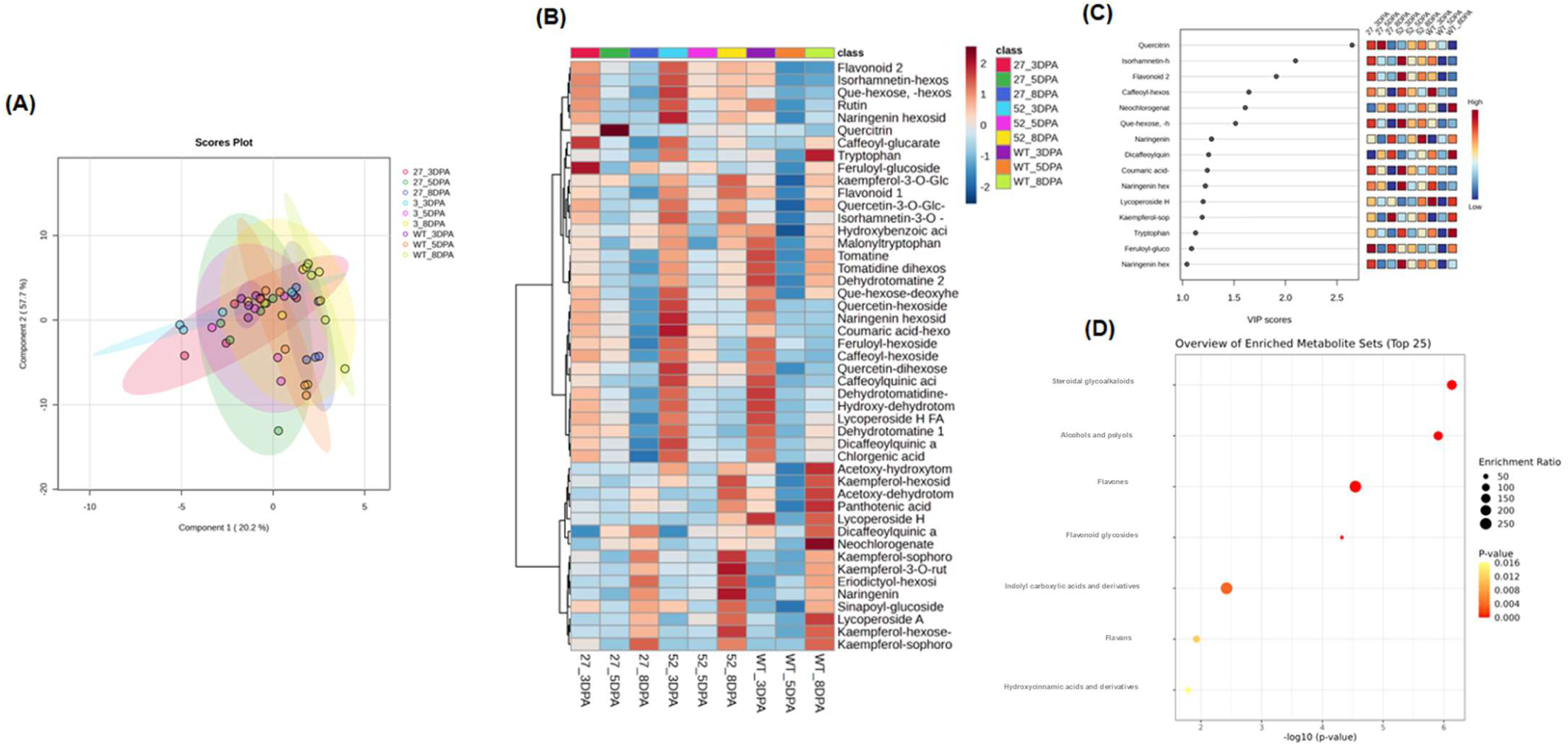
Metabolic characterization of tomato fruit at three developmental stages −3, −5, and 8 days post anthesis (DPA) collected and analyzed by LC–MS. (A) Partial least square-discriminant analysis (PLS-DA). (B) Heat map representation of the metabolite contents identified in this study. (C) The panel shows the top 15 variable importance in projection (VIP) identified by PLS-DA. The colored boxes on the right indicate the relative concentrations of each metabolite at each stage of development. (D) Quantitative enrichment analysis (QEA) overview presenting the top 25 related metabolic pathways ranked according to the P value. The data were normalized by using Log and Auto-scaling transformations on the MetaboAnalyst platform 6.0 (n = 4-6).

**Figure S8.**
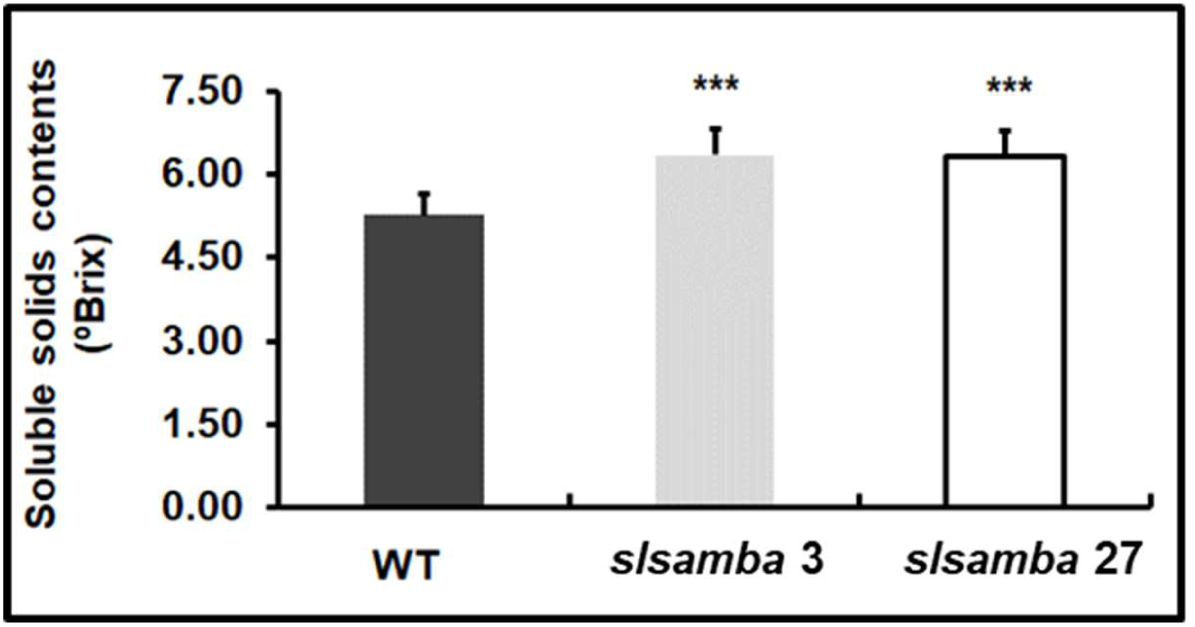
Soluble solids (°Brix) of red ripe tomato fruits from *slsamba* lines and wild-type plants. Values means ± SE.

**Figure S9.**
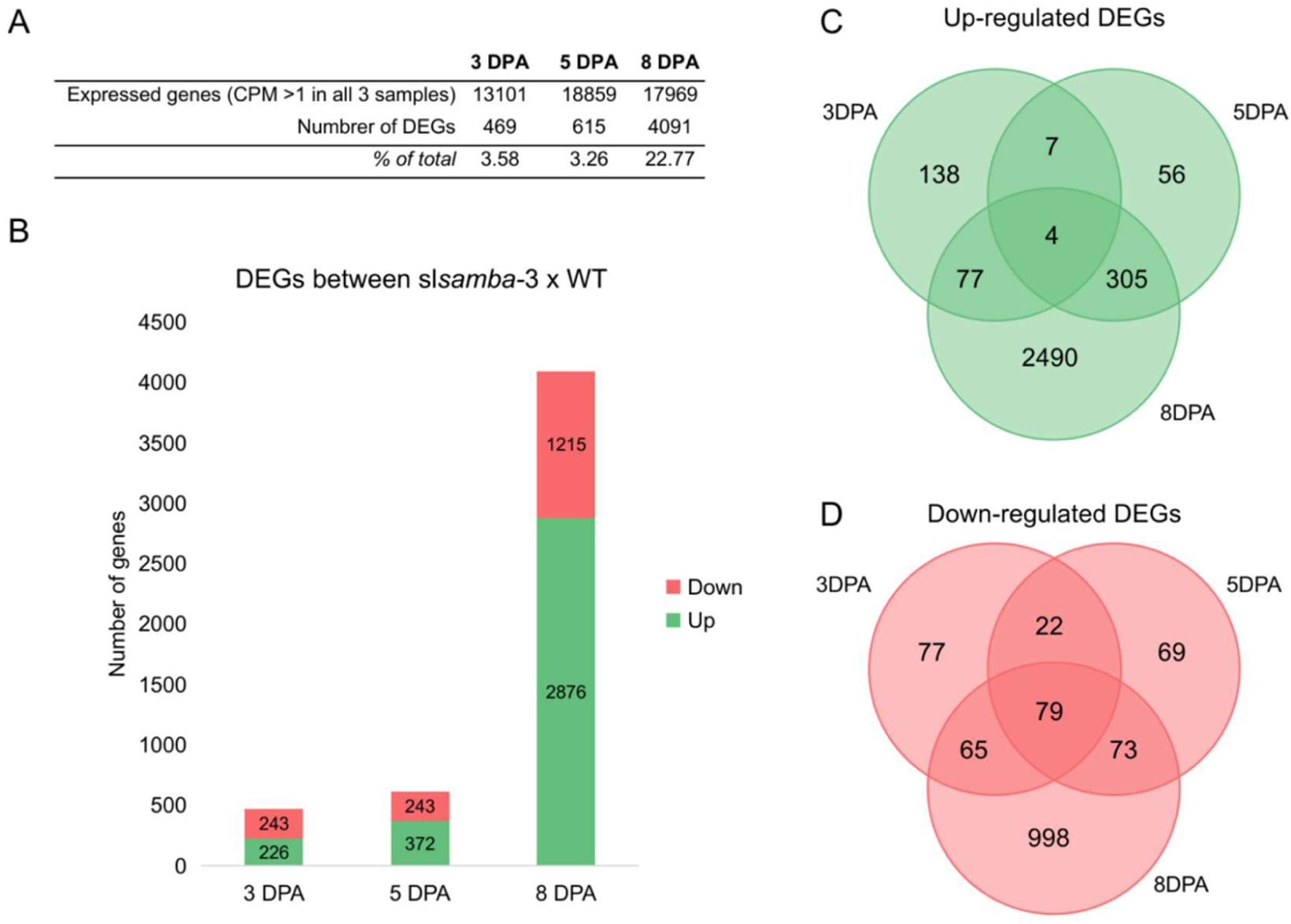
Results of differential expression analysis between *slsamba-3* and WT fruits. (A) Number of DEGs compared to the total expressed genes at each time point. (B) Bar plot of the number of DEGs at 3 DPA, 5 DPA, and 8 DPA, indicating up-regulated (green) and down-regulated (red) genes. (C) Venn diagram of up-regulated genes at all three time points. (D) Venn diagram of down-regulated genes at all three time points.

**Figure S10.**
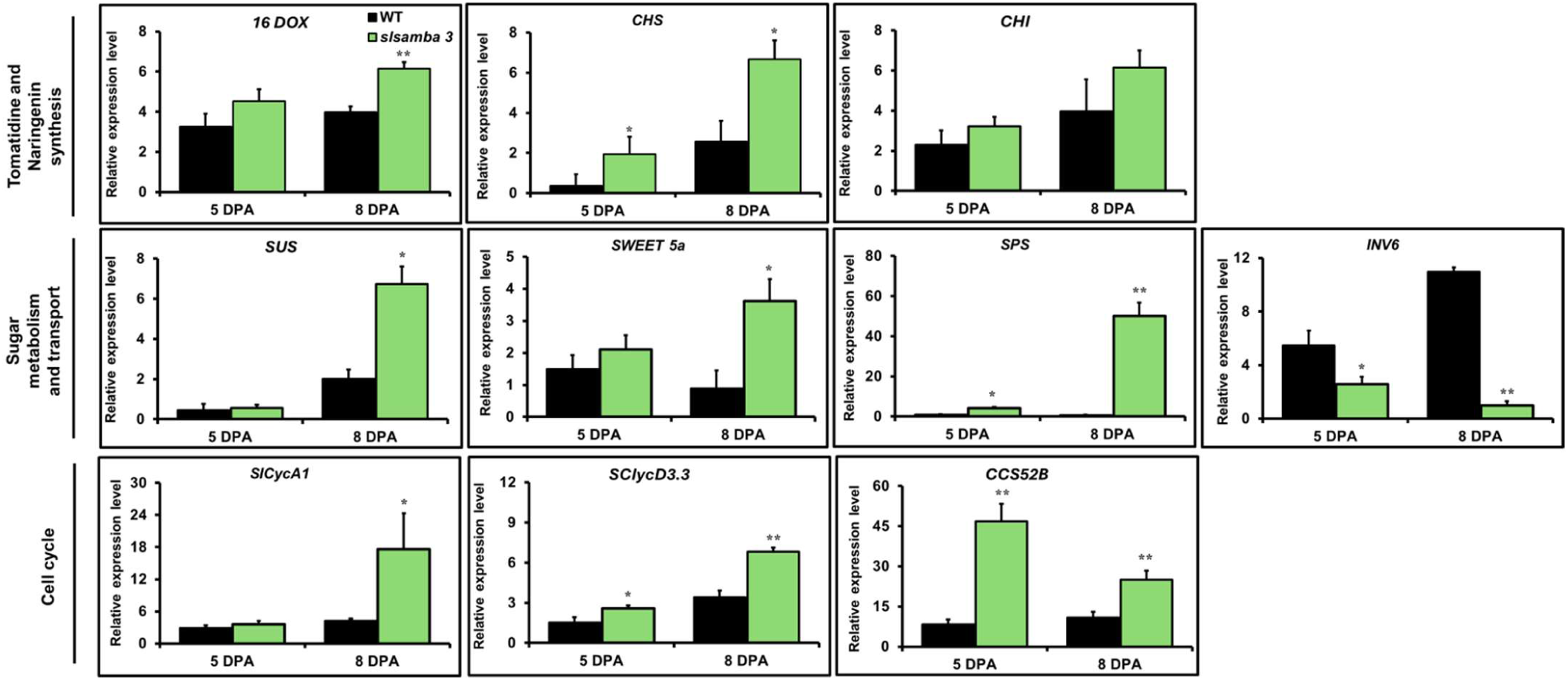
Relative expression by qRT-PCR in independent samples of *Solyc10g018190*1 (*16alpha,22,26- Trihydroxycholesterol* – *16DOX*), *Solyc01g090600* (*Chalcone synthase* – *CHS*), *Solyc05g010310* (*Chalcone-flavone isomerase* - *CHI*), *Solyc03g098290* (*Sucrose synthase* - *SUS*), *Solyc03g114200* (*SWEET 5a*), *Solyc08g042000* (*Sucrose phosphate synthase* - *SPS*), *Solyc10g083290* (*Invertase 6* – *INV6*), *SICycA1* (*Solyc11g005090*), *SlCycD3.3* (*Solyc04g078470*), *Solyc12g056490* (*CCS52B*) in line 3 and WT fruits harvested at three developmental stages (5-, 8- and 52-days post anthesis). Bars represent standard errors (SEs) of three biological replicates and two technical replicates. Significant differences (ANOVA followed by Dunnett’s t test) are shown by asterisks (*P < 0.05 and **P < 0.01).

**Table S1.**
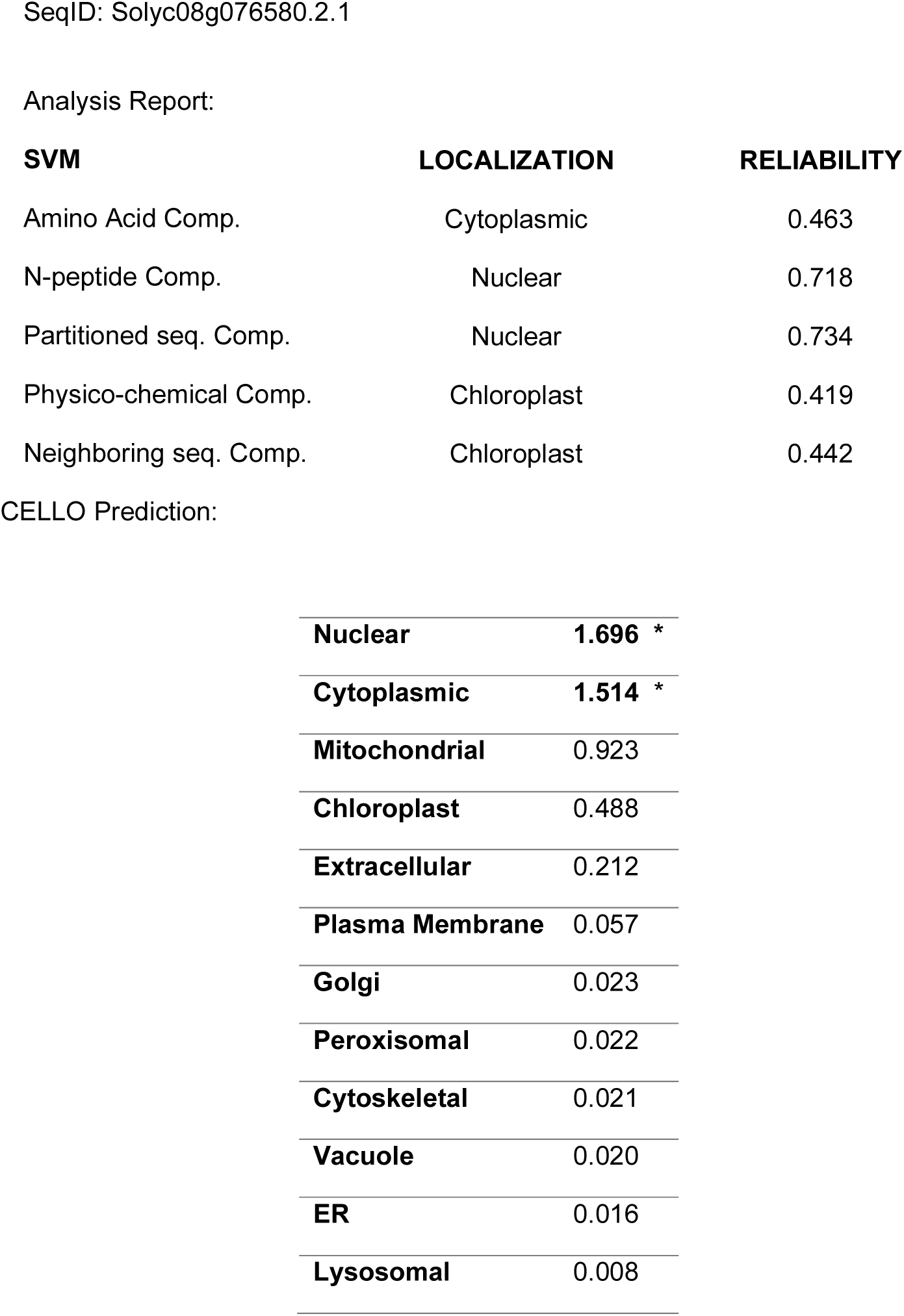
Prediction of SISAMBA subcellular localization by Cello.

| **SVM** | **LOCALIZATION** | **RELIABILITY** |
| --- | --- | --- |
| Amino Acid Comp. | Cytoplasmic | 0.463 |
| N-peptide Comp. | Nuclear | 0.718 |
| Partitioned seq. Comp. | Nuclear | 0.734 |
| Physico-chemical Comp. | Chloroplast | 0.419 |
| Neighboring seq. Comp. | Chloroplast | 0.442 |

|  |  |
| --- | --- |
| **Nuclear** | **1.696 \*** |
| **Cytoplasmic** | **1.514 \*** |
| **Mitochondrial** | 0.923 |
| **Chloroplast** | 0.488 |
| **Extracellular** | 0.212 |
| **Plasma Membrane** | 0.057 |
| **Golgi** | 0.023 |
| **Peroxisomal** | 0.022 |
| **Cytoskeletal** | 0.021 |
| **Vacuole** | 0.020 |
| **ER** | 0.016 |
| **Lysosomal** | 0.008 |

**Table S2.**
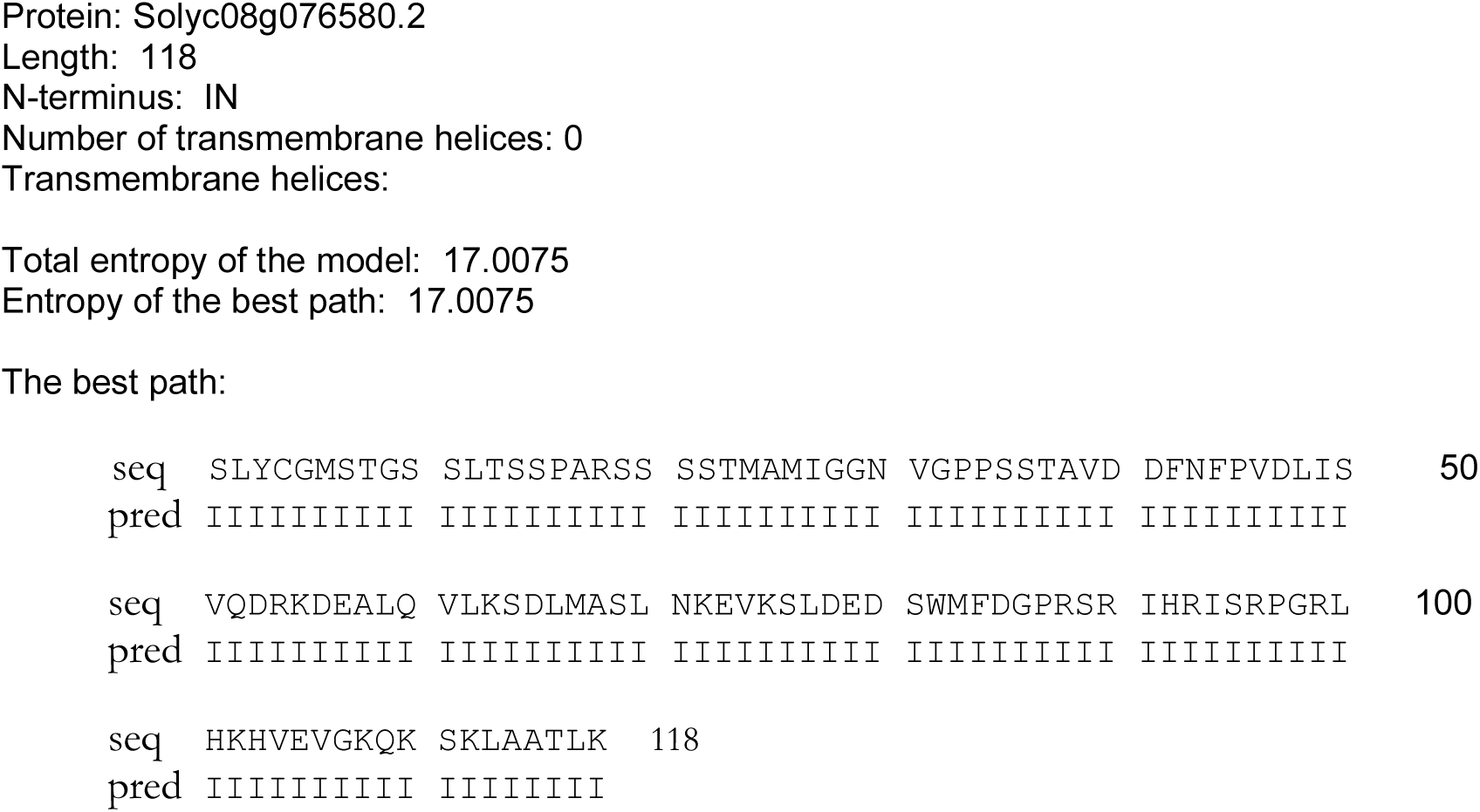
Transmembrane prediction of the Solyc08g076580.2 (SlSAMBA) protein through the HMMTOP.

**Table S3.**
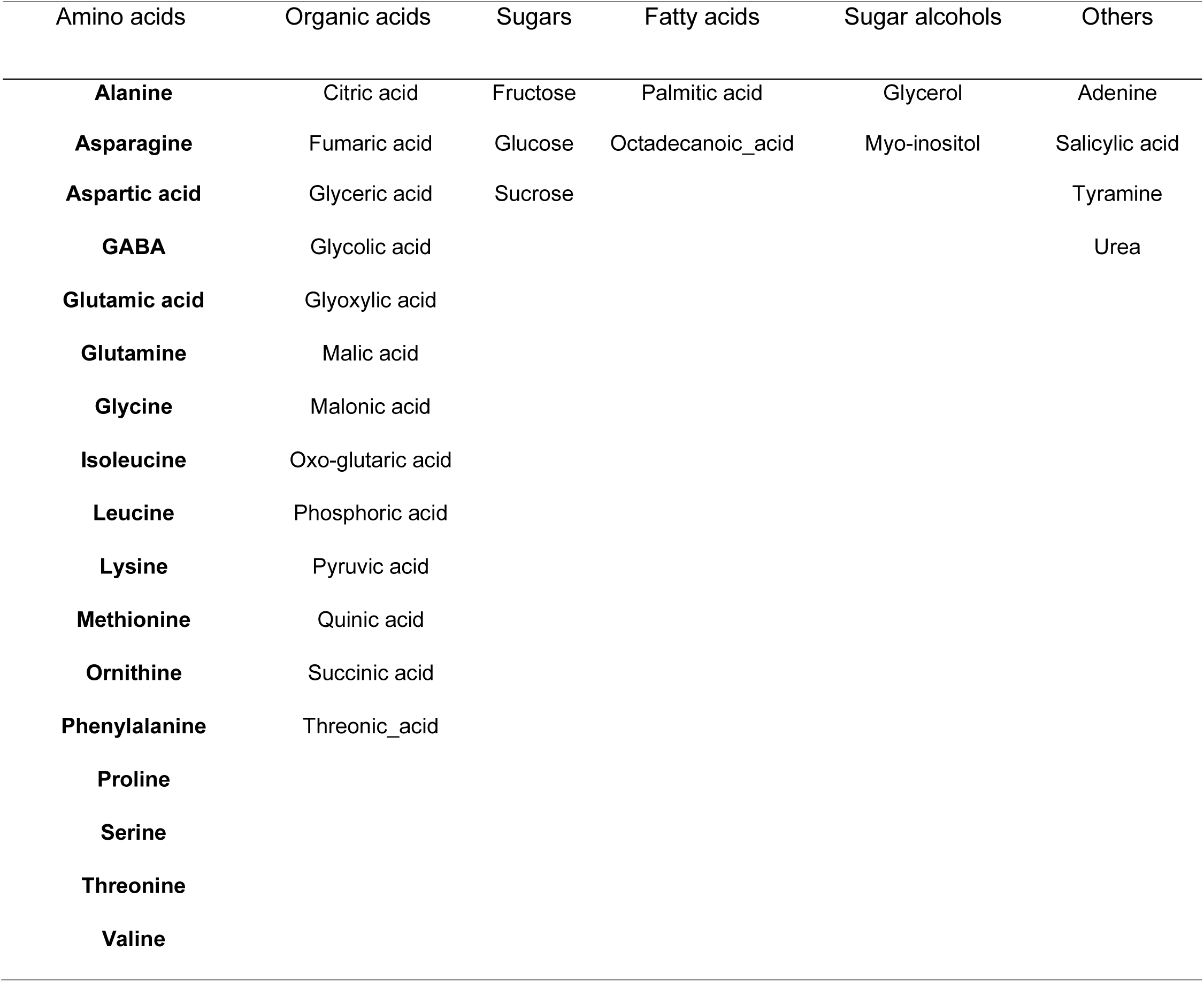
List of the 41 metabolites identified by GC–MS of fruits at −3, −5, or −8 days post anthesis (DPA) from *slamba* and WT lines.

**Table S4.**
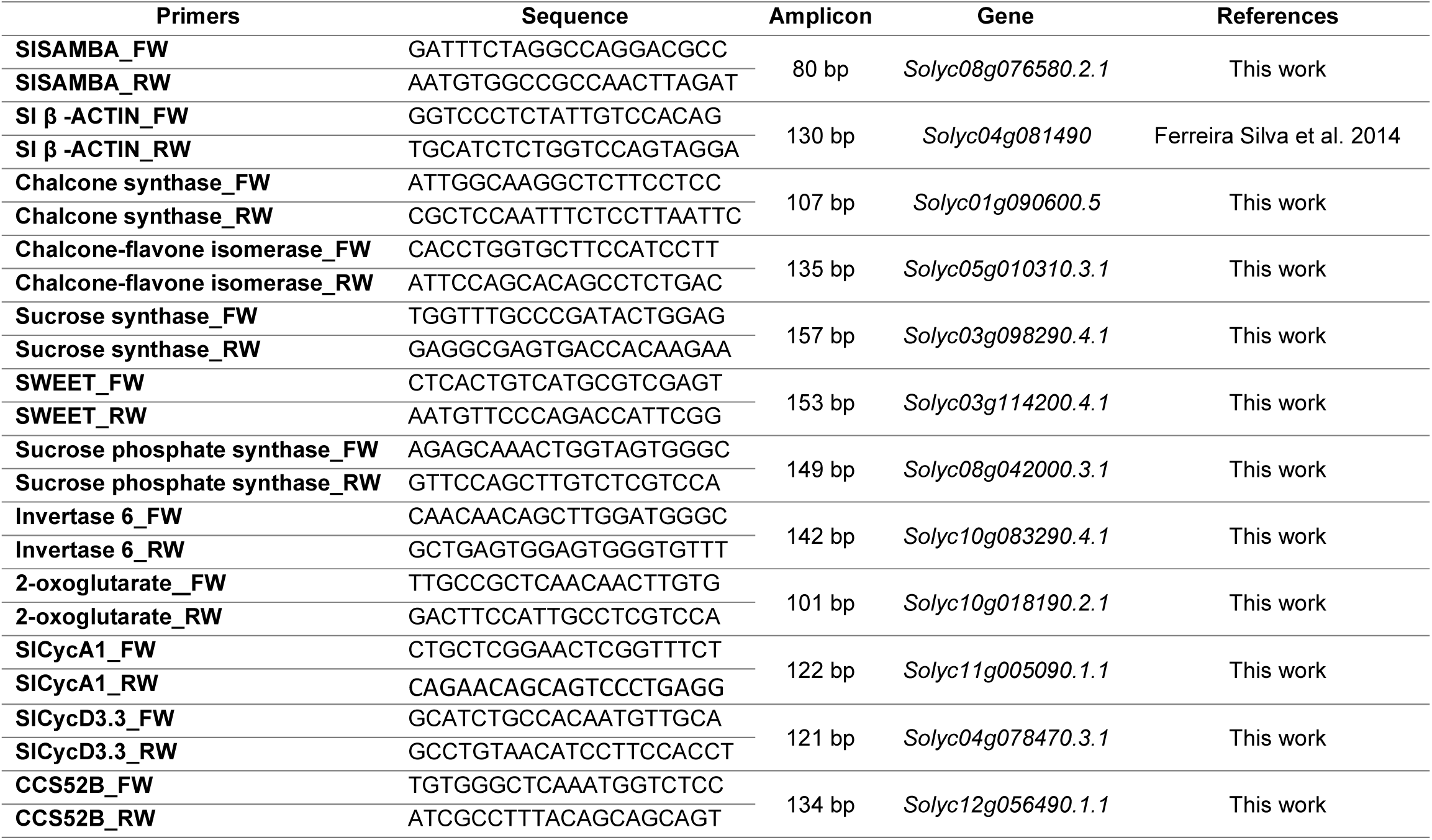
Oligonucleotide sequences used in this work.

